# Macro-level causal discovery from single-time-point observations of ecological dynamical systems

**DOI:** 10.1101/2024.10.10.608447

**Authors:** Daria Bystrova, Charles K. Assaad, Sara Si-moussi, Wilfried Thuiller

## Abstract

Ecologists often aim to uncover causal relations within ecological systems using observational data. However, many studies still rely mainly on correlation-based approaches, which do not support causal interpretation. Causal discovery methods have recently gained interest, particularly for ecological time-series data. Yet such data are often scarce, as collecting repeated and consistent measurements over long periods is costly, require strict continuation of protocols and is time-consuming. Consequently, ecologists have often access only to single-time-point observational data generated by dynamical systems. In this setting, unobserved past states can induce substantial unmeasured confounding, limiting the ability of standard algorithms such as PC and FCI to recover micro-level causal relations between observed variables. We show that PC and FCI can nevertheless recover meaningful causal information. In particular, they can identify specific cluster-level structures, which we call clustered super-unshielded colliders, which provide information about the partial causal ordering of macro-level variables. We further show that both algorithms can be reduced to a simple procedure, which we call RestPC, that yields the same identifiable information. We illustrate our results using simulated data and two real-world datasets: one on bird abundance, climate, and land cover, and another on soil microbial communities, environmental, terrain, and geochemical variables.

## 1 Introduction

Understanding the responses of ecological systems, particularly biodiversity, to changing environmental conditions is key to devising strategies for mitigating and adapting to ongoing land use and climate changes [90, 34]. It is also essential for establishing resilient protected areas [46]. Furthermore, due to the multispecies nature of many current disease threats, grasping these ecological dynamics is key to bolstering public health initiatives aimed at controlling infectious diseases [44, 83]. The distribution of species across space is theoretically contingent on abiotic factors (e.g., climate or soil conditions) as well as interactions among species. These interactions encompass various aspects such as food resources (e.g., predation, pollination), parasitism, and competition for space, habitat, or resources [38]. Despite this basic knowledge, unraveling the intricate relationships between the myriad species within diverse ecosystems under varying abiotic conditions poses significant challenges to the vast array of organisms and potential interaction types [91].

Ecological communities are represented by dynamic systems for which the composition and the structure changes over time in response to environmental conditions and biotic interactions. Consequently, ecological data are ideally collected as multiple time series (spacio-temporal data). Such data may be analyzed using methods such as Multivariate AutoRegressive models [42, 81] and Dynamic Bayesian Networks [8, 80], which can be used to infer aspects of community structure from repeated observations. It is important to note, however, that these methods are not causal by default. While they can capture temporal dependencies, they do not by themselves justify a causal interpretation unless additional assumptions are introduced explicitly. Indeed, temporal dependence may reflect direct causal effects, but also indirect pathways, hidden confounders, or shared responses to environmental variation. By contrast, methods that explicitly introduce assumptions to support causal discovery from time-series data [36, 37, 87, 89, 50, 79, 77, 4, 5] have recently gained increasing attention in ecology [11, 96, 78].

However, due to the complexity and cost of collecting data in such systems, it is challenging to obtain observations at multiple timepoints. As a result, some datasets often consist of a single time-point, capturing only a snapshot of the ongoing processes. For example, most available community data, derived from sources like GBIF, environmental DNA metabarcoding, and standardized in-situ protocols, often manifest as single-time-point observations rather than time-series (with multiple time steps). A figure illustrating the difference between single-time-point data and spatiotemporal data is provided in the Supplementary Material. Consequently, a comprehensive understanding of past states in ecological dynamic systems is often lacking.

Therefore, as in the time-series setting, several non-causal methods have been proposed to infer species interactions from non-temporal data. These include null-model approaches [35], species distribution models incorporating abiotic factors, such as joint species distribution models [71, 59], and methods that model statistical dependencies among ecosystem components, such as Markov-network-based approaches [56, 72, 20]. For comprehensive overviews, see [12, 70]. These methods have provided valuable insights into ecological systems. For example, they can identify nonrandom patterns of species co-occurrence, quantify associations between species and environmental variables, model residual dependence among taxa after accounting for abiotic factors, and describe conditional statistical dependencies among ecosystem components. However, their ecological interpretation remains challenging when the goal is to infer causal interactions. They are not designed to recover causal structure, nor do they necessarily recover the full conditional independence structure among ecosystem components. The ecological literature is not blind to this issue, and various arguments have been presented to elucidate the challenge of retrieving causal relations among ecosystem components from observational data [12, 70].

Bayesian networks [63, 53, 18] partly address this issue and can therefore distinguish some direct from indirect associations. In particular, unlike previous approaches, they can reveal that two species are independent once one conditions on a third variable, thereby reducing some spurious associations. However, such graphical structures are not causal by default: without additional assumptions, the directions of the edges cannot in general be interpreted as causal relations. Intriguingly, outside the ecological literature, substantial effort has been devoted to *causal discovery* from *non-temporal* observational data, that is, to identifying the assumptions under which causal relations can be recovered [65, 86, 33]. Among the best-known approaches in causal discovery from non-temporal data are constraint-based methods [86], which rely on conditional independence tests to recover causal relations. However, these methods are typically developed for data generated by systems that are not dynamic, or at least for settings in which temporal dependencies can be ignored.

This is a major limitation in ecology, where many variables are shaped by dynamic processes unfolding over time. As a result, learning causal relations from single-time-point data generated from a dynamic system is considerably more difficult than learning them from temporal data, which better capture changes in species abundances and distributions driven by environmental conditions, soil properties, and biotic interactions [12]. Indeed, such data may reflect a processes whose past states are unobserved, and these unobserved past states may act as unmeasured confounding of the variables observed at the present time, making causal discovery substantially more difficult. In such a setting, it is well known that micro-level causal relations cannot, in general, be discovered [36, 50, 79, 77]. This raises a fundamental question: although micro-level causal relations cannot generally be recovered from single-time-point observational ecological data, can causal discovery algorithms still recover meaningful process-level, or macro-level, causal information? Motivated by this question, our contributions are as follows: (i) we investigate which macro causal features remain identifiable, and which do not, from such data using constraint-based causal discovery algorithms [86], under assumptions that are either standard in the causal discovery literature or natural in ecological applications; (ii) we introduce RestPC, a new algorithm that simplifies existing methods and reduces computational time while maintaining the same results; (iii) we support our theoretical analysis with empirical tests on simulated data and illustrate the methods on real-world datasets.

The remainder of the paper is organized as follows: Section 2 introduces preliminaries. Section 3 investigates the applicability of causal discovery when we have single-time-point observational data. Section 4 validates our findings through simulation studies and Sections 5 and 6 illustrate them using two real datasets: one containing bird-species abundance records together with climatic and land-cover variables, and another containing environmental, terrain, geological and soil characteristics. Section 7 provides a discussion of the results. Proofs are deferred to the Supplementary Material and detailed code for the experiments can be found at https://github.com/CIPHOD/pyCIPHOD/tree/main/reproducibility/philosophicaltransactions2026/.

## 2 Preliminaries

In this section, we present pivotal concepts, tools, and assumptions. A directed acyclic graph (DAG) *G* is comprised of vertices denoted as V and edges as E. An edge links two vertices within the DAG *G* if they are directly causally linked^1^. In a dynamic system, the DAG is often referred to as a full-time DAG to emphasize that it represents the entire system and incorporates temporal information within the DAG. Throughout the paper, we will use the terms full-time DAG, and DAG interchangeably. Because we consider only a single-time-point, namely a snapshot of an underlying dynamic system, most vertices in the corresponding full-time DAG are unobserved. In the figures, unobserved vertices are represented in gray and observed vertices in white. Figure 1(a,e,i,m) illustrates several examples of such full-time DAGs. We note all vertices at time *t* in a full-time DAG as V_*t*_ and *t* is assumed to be sufficiently far from the lower boundary of the time domain. It is usually considered that such full-time DAG is compatible with a probability distribution over the variables (represented by vertices in the full-time DAG). This compatibility is given by the causal Markov condition [86].

**Figure 1.**
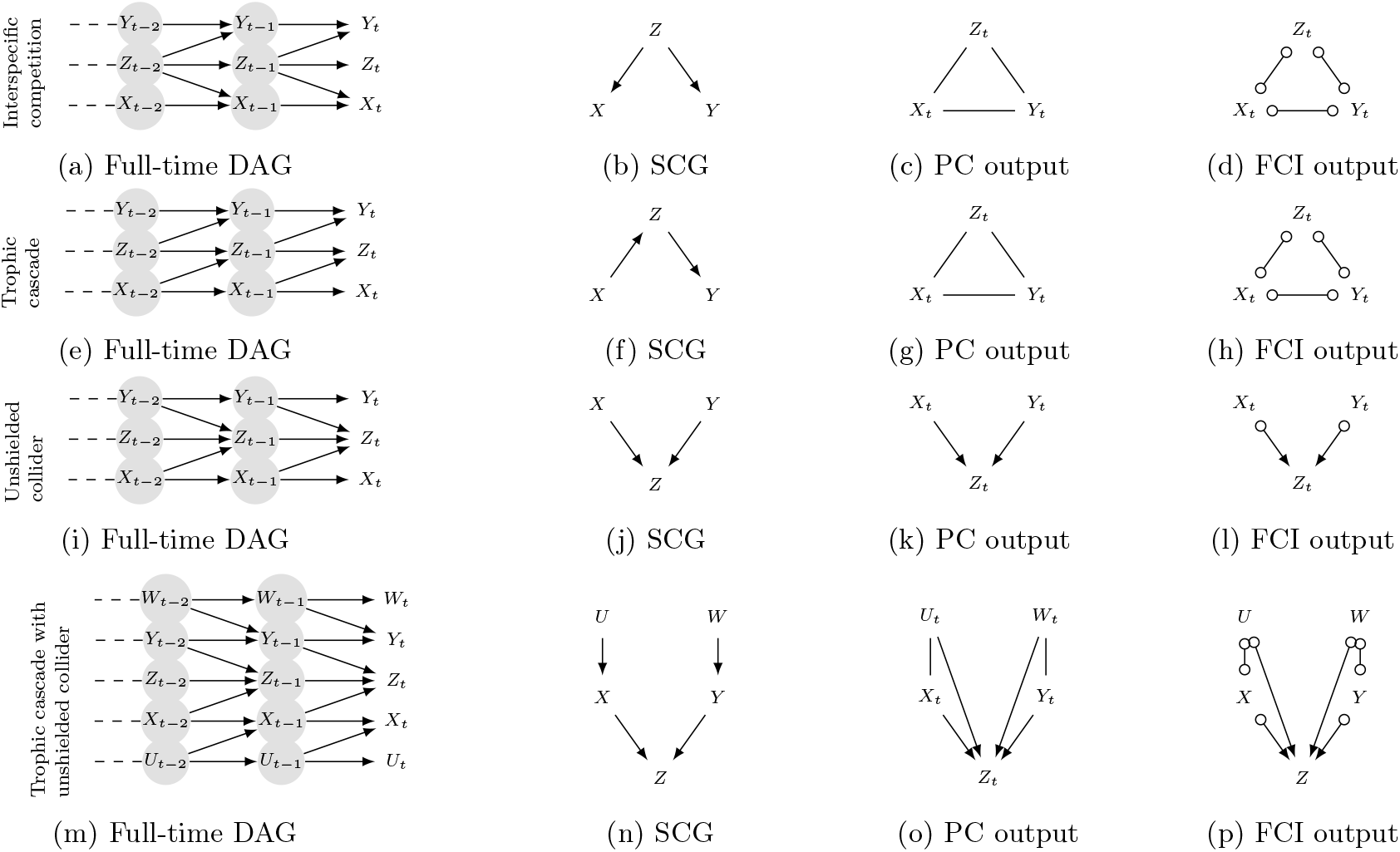
Illustration of full-time DAGs (a, e, i, m) with corresponding SCGs (b, f, j, n), CPDAGs and PAGs respectively inferred by PC (c, g, k, o) and FCI (d, h, l, p) from single-time-point observational data. Among the SCGs presented here, only the SCGs in (j) and (n) contain a CSUC. In (j), the CSUC is ({*X*}, {*Z*}, {*Y*}), while in (n), the CSUC is ({*U, X*}, {*Z*}, {*W, Y*}).

**Figure 2.**
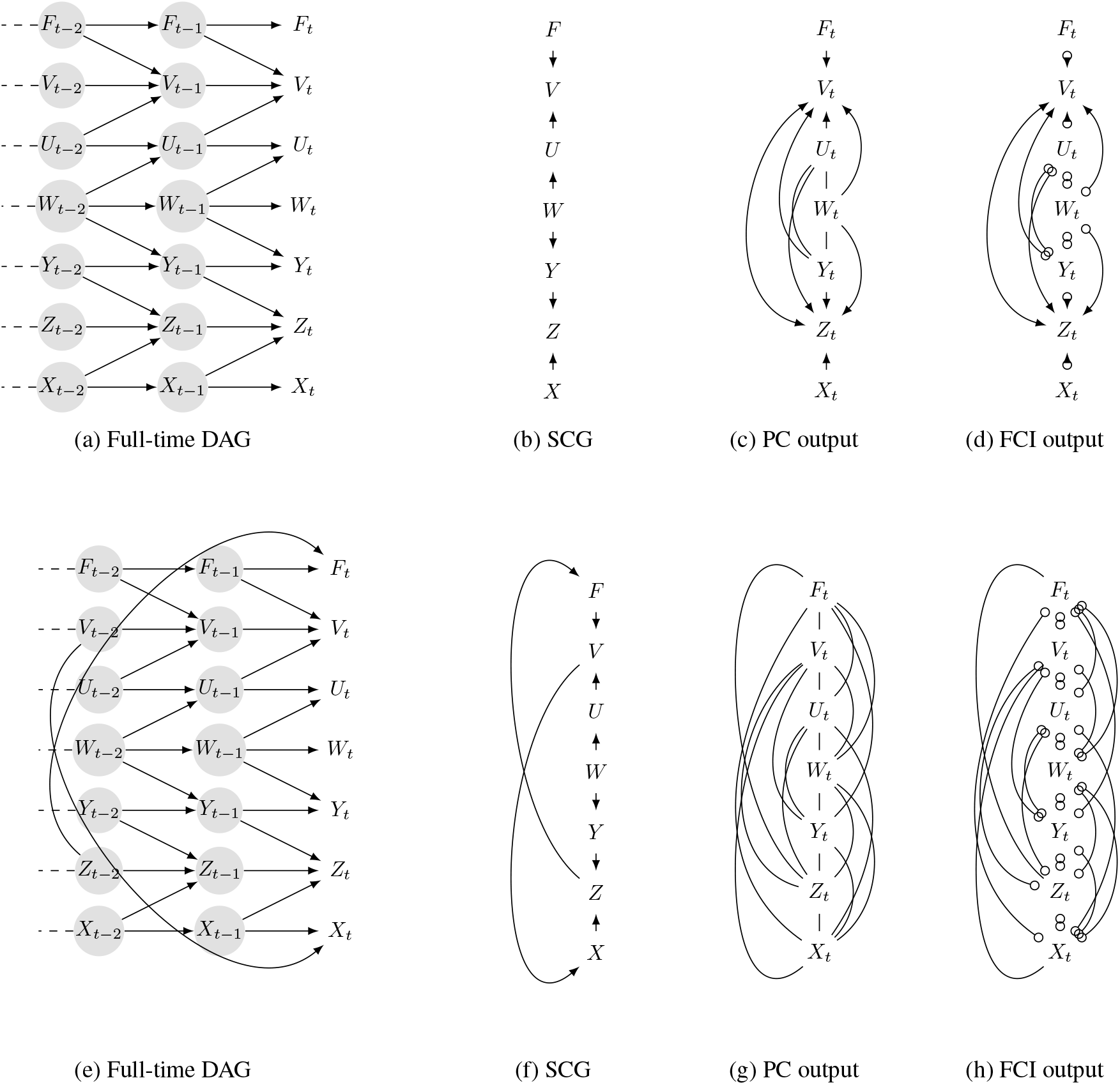
Illustration of more complex full-time DAGs (a,e) with corresponding SCGs(b,f), CPDAGs and PAGs respectively inferred by PC (c,g) and FCI (d,h) from corresponding single-time-point observational data. Among the SCGs presented here, only the SCGs in panel (b) contain CSUCs, namely({*F*}, {*V*}, {*U, W, Y, Z*}) and ({*V, U, W, Y*}, {*Z*}, {*X*}).

### Assumption 1

(Causal Markov condition). *Let G be a full-time DAG with a vertex set* V *and let P be a probability distribution over the vertices in*V. *For every Y*_*t*_ ∈V, *Y*_*t*_ *is independent in P of its non-descendants conditional on its parents in G*.

In simpler terms, a variable *W*_*t*_ is independent of all other variables (except those it influences) knowing (i.e., conditioned on) all its direct causes. For example, in a simple trophic chain where carnivores feed on herbivores and herbivores feed on plants (carnivores → herbivores → plants), plant abundance is independent of carnivore abundance when herbivore abundance is held constant. This causal Markov assumption is implicit in linear models such as regression analysis, path analysis or factor analysis.

Assumption 1 allows us to identify certain independencies in the distribution that is compatible with the full-time DAG. However, it turns out that there are many more conditional independencies that can be read off the full-time DAG but are not explicitly covered by Assumption 1. To identify all these conditional independencies in a DAG, we can use a tool called d-separation [64]. For a full-time DAG G, two vertices, *X*_*i*_ and *Y*_*t*_ are d-separated given a set ℤ, written *X*_*i*_ ╨ _*G*_*Y*_*t*_ | ℤ, if and only if ℤ blocks all paths between *X*_*i*_ and *Y*_*t*_. A path is *blocked* by ℤ if and only if at least one vertex *Z*_*j*_ ∈ ℤ is a common cause ← *Z*_*j*_ → or an intermediate cause → *Z*_*j*_ → on this path, or if all colliders → *Z*_*j*_ ← on the path and their descendants are not in ℤ. A path that is not blocked by ℤ is said to be activated by ℤ.

In dynamic systems, and specifically in the time-series and stochastic process literature [6, 7, 3, 29, 75], an abstraction of the full-time DAG, called the *summary causal graph* (SCG) and denoted by *G*^*s*^, is often considered as it enables a more focused and parsimonious analysis of the causal structure by representing relations at the level of processes rather than individual time-indexed variables. In an SCG, a vertex *X* represents the entire process, and an edge *X* → *Y* indicates that there is at least one time-point *X*_*i*_ in the process *X* and a time-point *Y*_*t*_ in the process *Y* such that the edge *X*_*i*_ → *Y*_*t*_ exists in the full-time DAG. Figure 1(b,f,j,n) illustrates several examples of SCGs, many of which correspond to well-documented species interactions [97].

In general, deriving causal relations only from observational data is typically unfeasible. Nevertheless, in numerous specific scenarios and under additional varying assumptions, crucial insights can be gleaned that facilitate the partial or complete acquisition of causal knowledge. Various methodological families are available for the discovery of causal relations from non-temporal observational data [33], each grounded in its own distinct set of assumptions. In this paper, we focus on one of the most established and theoretically grounded categories, known as the constraint-based family of methods. Constraint-based causal discovery methods use statistical conditional independence tests to discover causal relationships from data.

However, solely employing Assumption 1 is not sufficient to uncover causal relations through conditional independence tests. This can be due to potential presence of deterministic relations, or canceling pathways. Consider two species, *S*^1^ and *S*^2^, alongside a specific environmental factor, temperature (*T*). In this scenario, an increase in *T*_*t*_ corresponds with elevated abundances of both 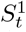 and 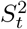. Yet, *S*^1^ preys on *S*^2^, leading to an indirect negative impact of *T*_*t*_ on 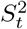. Despite the apparent causal relation between variables *T*_*t*_ and 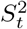, the opposing directions of direct and indirect effects, potentially canceling each other out. Consequently, this could result in an absence of dependence between *T*_*t*_ and 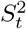 in the data, seemingly indicating independence that does not align with the DAG. Hence, the constraint-based causal discovery approach requires an additional assumption, called faithfulness, under which d-separation becomes equivalent to statistical conditional independence [86].

### Assumption 2

(Faithfulness, [86]). *A full-time DAG and a corresponding compatible probability distribution P are faithful to each other if, and only if, all the conditional independence relationships that hold true in P are implied by the Markov condition to G*.

This assumption is not testable from observational data alone and may be violated in ecological systems, particularly in the presence of stabilizing feedback mechanisms that maintain the system near equilibrium and mask underlying causal relations [1].

Furthermore, the presence of spurious correlations in the data due to unmeasured confounding constitutes an additional obstacle to the discovery of causal relations using conditional independence tests. Therefore, in conjunction with the previously mentioned assumptions, constraint-based approaches occasionally incorporate the **causal sufficiency over the DAG** assumption which states that there exist no unmeasured confounding between two observed variables. However, in ecology, confirming the absence of unmeasured confounders becomes intricate in complex ecosystems, where certain parts may remain unmeasured (e.g., unknown or difficult to obtain soil observations or measurements), rendering their influence on the observable part unknown.

Even under the above assumptions, recovering all causal relations from observational data remains infeasible, because several distinct DAGs may represent the same probability distribution [86]. However, these assumptions still make it possible to identify partial causal information, namely the features that are common to all DAGs that are Markov equivalent to the true DAG, that is, all DAGs implying the same d-separation relations (therefore implying the same conditional independencies). In particular, such DAGs share the same skeleton and the same unshielded colliders [94], namely collider configurations in which the two endpoints are not adjacent [86]. In ecology, unshielded colliders can arise when two distinct environmental factors influence the same ecological response. For example, consider Figure 1(i,j) and suppose *X* and *Y* are sulfur and nitrogen deposition respectively, and *Z* is the presence of plant species (see Figure 3(a) in [82]).

**Figure 3.**
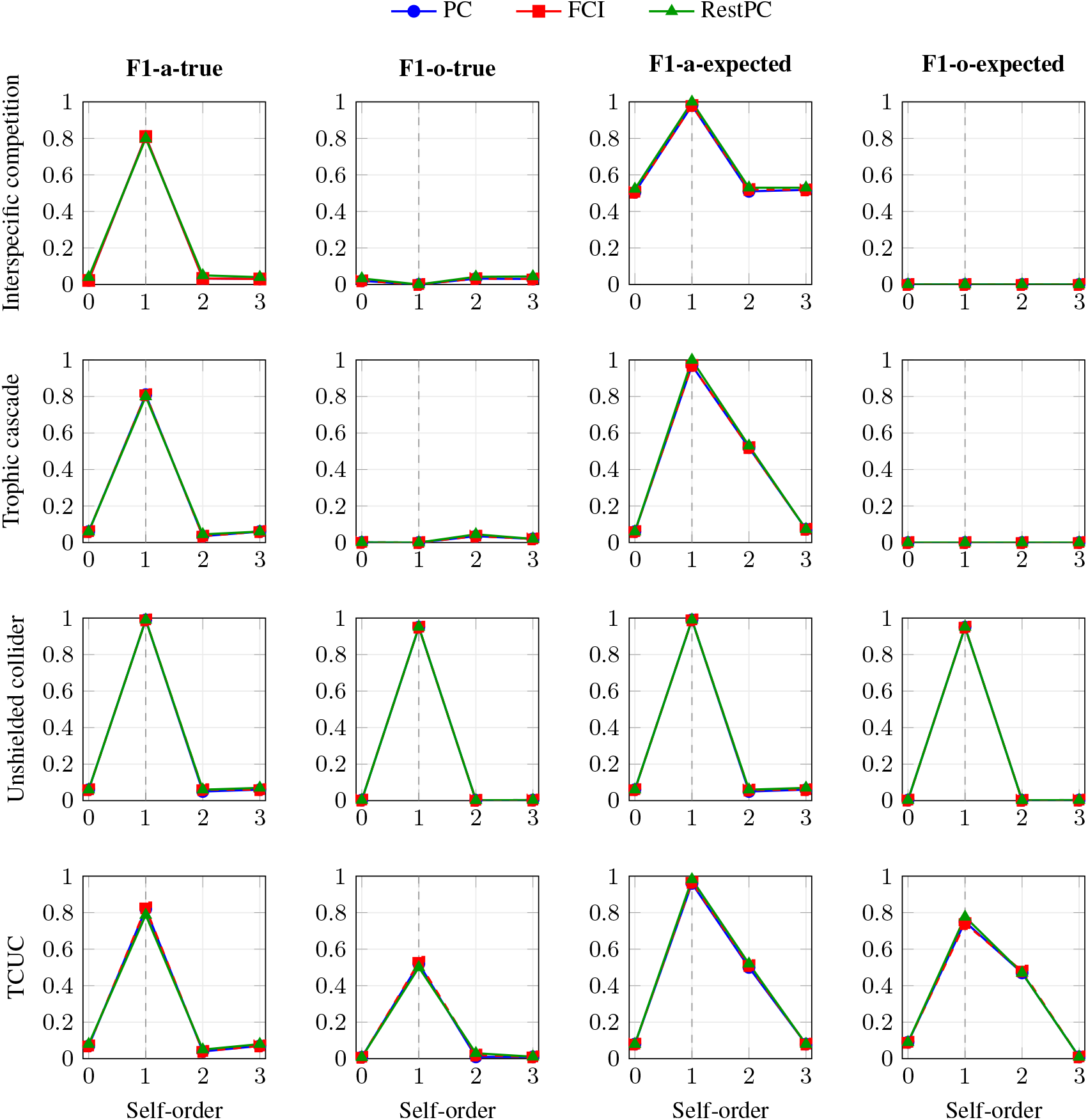
Sensitivity analysis to the self-dependence lag where TCUC is trophic cascade with unshielded collider. The lag of every cross-series causal relation is fixed to 1, while the self-dependence lag is varied from 0 to 3, with 0 denoting the absence of self-dependence. The dashed vertical line indicates the first-order self-dependence setting, which is the only setting satisfying the condition of Proposition 1. Performance is reported for PC, FCI, and RestPC across the different causal structures and graphical features considered.

The PC algorithm [86] is one of the most prominent constraint-based methods for causal discovery. Under Assumptions 1, 2, and **causal sufficiency over the DAG**, the PC algorithm does not, in general, identify a unique DAG. Instead, it recovers a CPDAG (Completed Partially Directed Acyclic Graph) [2, 19]. More precisely, the CPDAG contains the same skeleton as every DAG in the equivalence class, and it orients exactly those edges whose direction is shared by all DAGs in the class. Edges whose orientation differs across equivalent DAGs remain undirected. We briefly describe the main steps of the PC algorithm. Its first phase aims to recover the undirected skeleton of the underlying DAG. To do so, the algorithm proceeds as follows: **Step 1.1**. The algorithm starts from a fully connected undirected graph 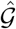, in which each vertex represents an observed variable (e.g., species abundance, a biodiversity indicator, or an environmental variable). **Step 1.2**. It removes from 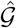 edges between pairs of vertices that are found to be marginally independent. **Step 1.3**. It then examines pairs of vertices that remain adjacent in 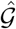 and removes the edge between them whenever they are found to be conditionally independent given some subset of vertices. The second phase of the PC algorithm aims to orient as many of the remaining edges as possible. It proceeds as follows: **Step 2.1**. The algorithm first identifies all unshielded colliders using the following rule: given an unshielded triple *X*_*t*_ − *Z*_*t*_ − *Y*_*t*_ in 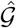, if *Z*_*t*_ was not part of the conditioning set that rendered *X*_*t*_ and *Y*_*t*_ independent in the first phase, then the triple is oriented as *X*_*t*_ → *Z*_*t*_ ←*Y*_*t*_. **Step 2.2**. Additional edges are then oriented using the Meek rules [52], which exploit the unshielded colliders identified in the previous step. For more details on the PC algorithm, see [86, 21].

An important extension of the PC algorithm is the FCI (Fast Causal inference) [86] algorithm, which relaxes the assumption of **causal sufficiency over the DAG**. This additional flexibility comes with a cost. Instead of returning a CPDAG, FCI outputs a PAG [76] (Partial Ancestral Graph), a more complex object that represents the causal features that remain identifiable despite unmeasured variables. PAGs are therefore more general, but also more difficult to interpret than CPDAGs. For instance, unlike CPDAGs, which contain only two types of edges (−, →), a PAG can contain up to five types of edges 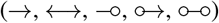. Given its greater complexity, and in the interest of keeping the main text clear and focused, a detailed presentation of the FCI algorithm is deferred to the Supplementary Material. Note that both the PC and the FCI algorithms allow incorporating prior knowledge by forbidding or forcing certain edges and by forbidding or forcing certain orientations [52]. This might be of particular interest in ecology, when for instance we have prior knowledge on who eats whom, or want to avoid having species abundance influencing global temperature. All the wealth of existing knowledge in ecology might thus be easily incorporated and would likely help the causal discovery algorithms. Furthermore, these methods can be applied on mixed data, a common feature of ecological datasets in which variables may include counts, presence–absence, biomass, or qualitative environmental descriptors, either through conditional independence tests for mixed data [22, 41, 98] or through preprocessing strategies such as discretization or nonparanormal transformations [49, 85].

In addition to Assumptions 1 and 2 required by constraint-based causal discovery algorithms, we will make additional three assumptions related to the dynamic system and the processes. The first of these assumptions is consistency throughout time.

### Assumption 3

(Consistency throughout time in direction). *A full-time DAG is said to be* consistent throughout time in direction *if all causal relations remain constant in direction throughout time. More precisely, if the causal strength associated with the edge* (*X*_*t*−*l*_ → *Y*_*t*_) *is nonzero for one admissible time point* (*t*), *then the corresponding causal strength associated with* (*X*_*s*−*l*_ → *Y*_*s*_) *is also nonzero for every admissible time point* (*s*).

This assumption implies that if *X*_*t*−1_ directly causes *X*_*t*_ in the full-time DAG, then *X*_*t*−2_ directly causes *X*_*t*−1_ as well. Crucially, however, it does not impose equality of the corresponding causal strengths. In contrast, many time-series causal discovery methods assume, in addition to Assumption 3, that causal strength remain constant over time [50, 79, 77, 6]. This stronger assumption is often unrealistic, for instance, in ecology, interaction strengths between species may vary over time due to changes in demography, population densities, or environmental conditions [23, 66, 55]. Our framework does not require such an assumption.

The second assumption we require is first-order self-dependence of each process, which is natural in many ecological dynamical systems [43, 40, 9], especially when data are collected yearly [13].

### Assumption 4

(First-order self-dependence). *For every process X and every time-point t, the state X*_*t*_ *is directly causally affected by its immediately preceding state X*_*t*−1_.

Under Assumptions 3 and 4, the assumption of **causal sufficiency over the DAG**, which is required by the PC algorithm, is rarely satisfied when dealing with single-time-point data. This is because the past will almost always contain unmeasured confounders. As a result, we replace this assumption with a weaker one, asserting causal sufficiency over the processes of the ecological system. Essentially, this means we must ensure that data is collected on every species or environmental variable that influences more than one species of interest.

### Assumption 5

(Causal sufficiency over the SCG). *For any two distinct observed variables X*_*t*_ *and Y*_*t*_, *arising respectively from processes X and Y, every process Z that causally affects both X and Y in the SCG is also observed at time t*.

For example, suppose that *X*_*t*_ and *Y*_*t*_ are observed variables and that the SCG contains (*X* ←*Z* → *Y*). This means that the process *Z* causally affects both processes *X* and *Y*. Under Assumption 5, *Z*_*t*_ must also be observed whenever *X*_*t*_ and *Y*_*t*_ are observed. Thus, the assumption rules out the case where *X*_*t*_ and *Y*_*t*_ are both observed while *Z*_*t*_ is unobserved. However, unlike **causal sufficiency over the DAG**, it does not require the past variables of *X*_*t*_, *Y*_*t*_, and *Z*_*t*_ to be observed. This distinction is important for the setting considered in this paper, where these past variables are explicitly supposed to be unobserved.

## 3 Discovering macro causal relations from single-time-point observational data

Micro-level causal relations cannot, in general, be identified from single-time-point observational data under our assumptions, a limitation already established in the literature [36, 50, 79, 77]. We illustrate this point with examples in the Supplementary Material. In this section, we investigate when *macro-level* causal relations, namely relations between the underlying ecological processes, can nevertheless be identified from such data. In particular, we study what the PC and FCI algorithms can reveal about causal relations at the level of processes. For this purpose, after applying either algorithm to the variables observed at time *t*, we introduce an additional step to derive an inferred SCG. More precisely, let 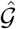 be the output of PC. We define the inferred SCG, denoted 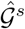, as follows. For every pair of observed variables *X*_*t*_, *Y*_*t*_ ∈V_*t*_, we include the edge *X* → *Y* in 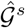 if and only if *X*_*t*_ → *Y*_*t*_ is present in 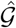, and we include the edge *X* − *Y* in 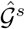 if and only if *X*_*t*_ − *Y*_*t*_ is present in 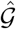. The inferred SCG for FCI is constructed analogously, with all endpoint marks preserved to reflect edge uncertainty in 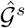.

If two processes *X* and *Y* belong to different connected components of the SCG, then the absence of a relation between them can be recovered, since in that case *X*_*t*_╨_*G*_*Y*_*t*_ | ∅. In the following, we therefore focus on the more challenging case in which *X* and *Y* belong to the same component of the SCG. Let us first consider two processes *X* and *Y* such that *X* → *Y* in the true SCG. In this situation, both PC and FCI will detect a connection between *X* and *Y*, because 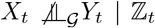, ∀ℤ_*t*_ ⊆V_*t*_ \ {*X*_*t*_, *Y*_*t*_}. This dependence may arise either from a direct causal relation between *X*_*t*_ and *Y*_*t*_, or from unmeasured confounding induced by the unobserved past. Consequently, although the existence of a relation between the two processes can be detected, its orientation cannot in general be recovered. In such cases, PC returns an undirected edge – and FCI returns an edge with circle marks (∘−∘).

Now consider three processes *X, Y*, and *Z*, in the setting of Figure 1(a), where *Z*_*t*−1_ acts as an unobserved confounding between *X*_*t*_ and *Y*_*t*_. The corresponding SCG is shown in Figure 1(b), where *Z* is a common cause of both *X* and *Y*. However, the output of the PC algorithm, shown in Figure 1(c), is an undirected triangle on *X*_*t*_, *Y*_*t*_, and *Z*_*t*_, and the SCG deduced from it is therefore also an undirected triangle. The reason is that *X*_*t*_ and *Y*_*t*_ remain dependent both marginally and conditionally on *Z*_*t*_, because they share unobserved common ancestors such as *Z*_*t*−1_, *Z*_*t*−2_, …. Conditioning on *Z*_*t*_, which is the only observed variable from process *Z*, cannot block the active paths linking *X*_*t*_ and *Y*_*t*_ through these past ancestors. A similar phenomenon occurs for the chain structure shown in Figure 1(e), whose true SCG is displayed in Figure 1(f). In that case as well, the PC algorithm returns a fully connected undirected graph (Figure 1(g)). Likewise, in these examples, FCI yields a fully connected graph with edges with circle marks (Figure 1(d,h)). In both examples, the inferred graph is therefore denser than the true SCG and it is unoriented.

The situation is different in the configuration shown in Figure 1(i), where *X*_*t*−1_ and *Y*_*t*−1_ both cause *Z*_*t*_. In this case, the outputs of PC and FCI, shown respectively in Figure 1(k) and Figure 1(l), lead to the correct SCG. Indeed, there is no active path between *X*_*t*_ and *Y*_*t*_, so that *X*_*t*_ ╨ _*G*_*Y*_*t*_ | ∅. Accordingly, both algorithms remove the edge between *X*_*t*_ and *Y*_*t*_. The triple (*X*_*t*_, → *Z*_*t*_, ←*Y*_*t*_) then forms an unshielded triple in the inferred graph, and since *Z*_*t*_ does not belong to the set that rendered *X*_*t*_ and *Y*_*t*_ conditionally independent, it is oriented as *X*_*t*_ → *Z*_*t*_ ← *Y*_*t*_. An analogous conclusion is obtained with FCI. Importantly, even though the configuration *X*_*t*_ → *Z*_*t*_ ← *Y*_*t*_ is not literally present in the full-time DAG, the SCG deduced from this orientation is *X* → *Z* ← *Y*, which is correct.

If in the preceding example, we imagine that there is a fourth process *W* such that *W* is a common cause of *X* and *Y* in the SCG, then 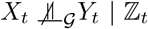, ∀ℤ_*t*_ ⊆V_*t*_ leading the PC and FCI algorithms to be unable to eliminate the edge between them. Consequently, we observe that the PC and FCI algorithms are only capable of identifying a subset of unshielded-colliders wherein no active path connects the endpoint vertices of the triples (such as *X* and *Y* in our example). Expanding on the previous example, if we introduce two more processes, *U* and *W*, where *U*_*t* − 1_ causes *X*_*t*_ and *W*_*t*−1_ causes *Y*_*t*_, as shown in Figure 1(m), the true SCG is illustrated in Figure 1(n). In this situation, both the PC and FCI algorithms can still orient the triple *X*_*t*_, *Z*_*t*_, *Y*_*t*_ correctly as *X*_*t*_ → *Z*_*t*_ ←*Y*_*t*_ and 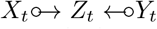 respectively, utilizing the same rationale as before. So in the deduced SCG, we get *X* → *Z* ←*Y*. However, both algorithms would incorrectly infer that both triples (*U, X, Z*) and (*W, Y, Z*) form a triangle. The reason is that *U*_*t*_ and *Z*_*t*_ and *W*_*t*_ and *Z*_*t*_ share common ancestors *U*_*t*−2_ and *W*_*t*−2_ (similar to the example in Figure 1(e)). Additionally, the PC and FCI algorithms would respectively orient the triple (*U*_*t*_, *Z*_*t*_, *W*_*t*_) as *U*_*t*_ → *Z*_*t*_ ← *W*_*t*_ and 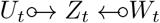. Which means in the deduced SCG we would have *U* → *Z* ← *W*, which is incorrect. Consequently, while PC and FCI correctly identify all unshielded colliders for which the extremities are marginally independent, they may identify additional unshielded colliders that are not true unshielded colliders. To address this issue and better characterize the structures that the PC and FCI algorithms can accurately detect, we introduce the concept of “clustered-super-unshielded-colliders”.

### Definition 1

(Clustered super-unshielded collider (CSUC)). *In an SCG G*^*s*^, *a* clustered super-unshielded collider (CSUC) *is a triple of nonempty, pairwise disjoint clusters of vertices* (X, ℤ, Y) *such that the following hold: 1*. ***Separation across sides:*** *for all X* ∈ X *and Y* ∈ Y, *there exists no active path between X and Y in G*^*s*^. *2*. ***Connection to the middle:*** *for all X* ∈ X *and Z* ∈ ℤ, *there exists an active path between X and Z in* ^*s*^ *such that all vertices on the paths are either in* X *or in* ℤ; *and for all Y* ∈ Y *and Z* ∈ ℤ, *there exists an active path between Y and Z in G*^*s*^ *such that all vertices on the paths are either in* Y *or in* ℤ. *3*. ***Maximality:*** *the triple* (X, ℤ, Y) *is maximal w*.*r*.*t. set inclusion among all triples satisfying (i)–(ii)*.

Although CSUCs are less immediately interpretable than standard unshielded colliders, they remain informative because they capture causal structure at the level of processes rather than individual variables. **A CSUC suggests that any process** *Z* ∈?ℤ **cannot cause any process** *X* ∈ X **or any process** *Y* ∈ Y, **since otherwise there would be an active path** *X* ←*Z* ←*Y* **or** *X* ←*Z* → *Y* **or** *X* → *Z* → *Y* **which are by definition not allowed. Hence, each process in** ℤ **cannot be causally upstream of any process in** X **and any process in** Y. Moreover, every pair (*X, Y*), with *X* ∈ X and *Y* ∈ Y, does not necessarily forms an unshielded collider with every process *Z* ∈ ℤ. However, for any CSUC (X, ℤ, Y), there exists at least one unshielded collider *X* → *Z* ←*Y*. This means that whenever the size of X, the size of ℤ and the size of Y are equal to 1, then the CSUC is an unshielded-collider. This is useful because, in our setting, single-time-point data generally do not allow reliable identification of micro-level causal relations between variables observed at time *t*. By contrast, CSUCs still provide interpretable information about the broader causal organization of the processes. Interestingly, the PC algorithm and the FCI algorithm are able to detect all CSUCs.

### Theorem 1.

*Consider a distribution satisfying Assumptions 1, 2, 3, 4, and 5 and suppose we are given perfect conditional independence information about all pairs of variables* (*X*_*t*_, *Y*_*t*_) *in*V_*t*_. *Then the PC and FCI algorithms recover the components of the SCG and the same process-level orientation information. In particular, both algorithms recover the process-level orientation information represented by the collection of CSUCs*.

Notice that to identify CSUCs in the setting considered in this paper, many steps of the PC algorithm are not needed. In particular, only the part of the algorithm needed to detect marginal independencies and identify CSUCs must be retained. Motivated by this observation, and to improve computational efficiency, we introduce a restricted version of the PC algorithm, called *RestPC*, that is adapted to this setting. In particular, RestPC retains only Steps 1.1, 1.2, and 2.1 of the PC algorithm.

### Corollary 1.

*Consider a distribution satisfying Assumptions 1, 2, 3, 4, and 5 and suppose we are given perfect marginal independence information about all pairs of variables* (*X*_*t*_, *Y*_*t*_) *in*V_*t*_. *Then the RestPC algorithm recovers the components of the SCG and the process-level orientation information represented by the collection of CSUCs*.

These results show that, at the population level, PC, FCI, and RestPC recover all CSUCs from single-time-point observational data. The advantage of RestPC is therefore not in recovering additional information, but in recovering the same information more efficiently. For example, RestPC requires only *d*(*d* − 1) marginal independence tests, whereas PC may require up to 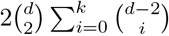 conditional independence tests in the worst case, when conditioning sets of size up to *k* are considered. Moreover, in practice, independence information must be estimated from finite data, and errors in independence testing may lead to spurious or missed CSUCs. Since RestPC relies on marginal rather than conditional independence tests, it may offer advantages [84] beyond computational efficiency, particularly in settings with limited sample size.

## 4 Simulation based on a known structure

In this section, we evaluate our theoretical findings using simulated single-time-point data. We consider four simple structural scenarios depicted in Figure 1: an interspecific competition, a trophic cascade, an unshielded collider, and a trophic cascade with unshielded collider. In each case, we simulate data using the following generator: 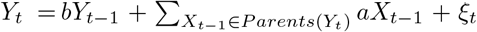. Here, *ξ*_*t*_ ~ *N* (0, 1), *b, a* ~ *U* (−1, 0.5) ∪ *U* (0.5, 1). For each run, we generate 5000 independent spatiotemporal series and retain only the final time point of each series, yielding a cross-sectional sample of size 5000. This entire procedure is then repeated 1000 times for each structural scenario, allowing us to estimate the mean and variance of the performance measures across runs.

Using the data generated from these simulations, we evaluate the PC, FCI, and RestPC algorithms using the Fisherz test [30, 47] while significance level *α* for the (conditional) independence tests is set to 0.05. To evaluate the performance of each method on simulated data, we consider four metrics. The first two compare the inferred graph with the true SCG: *F1-a-true*, the F1-score for adjacencies, and *F1-o-true*, the F1-score for orientations. The other two compare the inferred graph with the expected SCG: *F1-a-expected*, the F1-score for adjacencies, and *F1-o-expected*, the F1-score for orientations.

Our results are presented in Table 1. Across the four simulated scenarios, PC, FCI, and RestPC yield nearly identical results, which is expected in this setting, as discussed above. For interspecific competition and the simple trophic cascade, all methods recover the adjacency structure reasonably well (*F*1-a-true around 0.8), with substantially higher scores when evaluated against the expected structure (*F*1-a-expected around 0.98). By contrast, orientation with respect to the true graph is essentially null, reflecting the fact that, from a single observed time-point, these structures are not identifiable. In contrast, scenarios containing CSUCs are much better recovered. In unshielded collider, both adjacency and orientation scores are high, around 0.99 and 0.95, respectively. In the trophic cascade with unshielded collider, orientation recovery remains only partial when evaluated against the true graph (*F*1-o-true around 0.5), although adjacency recovery remains stable. However, when evaluated against the expected structure, performance is substantially higher, with adjacency and orientation scores around 0.97 and 0.76, respectively. Taken together, these results support our theoretical findings. In particular, for structures that do not contain CSUCs, the inferred graphs tend to be closer to fully connected graphs. At the same time, all methods recover CSUCs reasonably well.

**Table 1:** Mean F1-score and variance for detecting adjacencies and orientations using PC, FCI, and RestPC. F1-o-expected is reported as “NA” in the cases where the expected SCG contains no orientable structure.

|  |  | F1-a-true | F1-o-true | F1-a-expected | F1-o-expected |
| --- | --- | --- | --- | --- | --- |
| Interspecific competition | PC | $0.81 \pm 0.01$ | $0.0 \pm 0.0$ | $0.98 \pm 0.01$ | NA |
| | FCI | $0.81 \pm 0.01$ | $0.0 \pm 0.0$ | $0.98 \pm 0.01$ | NA |
| | RestPC | $0.81 \pm 0.01$ | $0.0 \pm 0.0$ | $1.0 \pm 0.0$ | NA |
| Trophic cascade | PC | $0.82 \pm 0.01$ | $0.01 \pm 0.01$ | $0.98 \pm 0.01$ | NA |
| | FCI | $0.82 \pm 0.01$ | $0.01 \pm 0.01$ | $0.98 \pm 0.01$ | NA |
| | RestPC | $0.8 \pm 0.01$ | $0.01 \pm 0.01$ | $0.99 \pm 0.01$ | NA |
| Unshielded collider | PC | $0.99 \pm 0.01$ | $0.95 \pm 0.04$ | $0.99 \pm 0.01$ | $0.95 \pm 0.04$ |
| | FCI | $0.99 \pm 0.01$ | $0.95 \pm 0.04$ | $0.99 \pm 0.01$ | $0.95 \pm 0.04$ |
| | RestPC | $0.99 \pm 0.01$ | $0.95 \pm 0.04$ | $0.99 \pm 0.01$ | $0.95 \pm 0.04$ |
| Trophic cascade with unshielded collider | PC | $0.82 \pm 0.01$ | $0.51 \pm 0.01$ | $0.97 \pm 0.01$ | $0.75 \pm 0.01$ |
| | FCI | $0.82 \pm 0.01$ | $0.51 \pm 0.01$ | $0.97 \pm 0.01$ | $0.76 \pm 0.01$ |
| | RestPC | $0.78 \pm 0.01$ | $0.5 \pm 0.01$ | $0.98 \pm 0.01$ | $0.78 \pm 0.01$ |

Additional settings involving feedback and nonlinear relations are reported in the Supplementary Material. We note that the nonlinear setting yields stronger contrasts between methods (RestPC clearly outperforms PC and FCI). This is likely because nonlinearity makes conditional independence testing substantially more challenging than marginal independence testing. In addition, the Supplementary Material include an empirical evaluation of the computation time of the three algorithms, showing, unsurprisingly, that RestPC becomes faster than the competing methods as the number of variables increases.

## 5 Structural relations between bird species and environmental variables

We stress at the outset that the analyses below are illustrative rather than validating: in the absence of independent ground truth, they characterize what PC and RestPC recover and how interpretable the results are, not which method is correct. Here we focused on data from the French Breeding Bird Survey (STOC), combining bird abundance records with environmental covariates describing climate and land cover [31]. We focus on the 30 most abundant bird species and 6 environmental variables, using data from 824 sites for which the last two consecutive years are available. See more details in the Supplementary Material.

Since our aim is to study what can be learned from single-time-point observations, the main analyses use only the most recent time point. We also consider the last two time points as a reference comparison based on additional temporal information. For the two-time-point setting, we learned a temporal graph using a temporal adaptation of PC (tPC [86]) in which we incorporate the background knowledge that environmental variables can not be the effects of the species abundances. However, the results obtained with tPC should not be considered ground truth, since tPC itself relies on causal discovery assumptions. Further details on the survey design, preprocessing, variable selection, and environmental data sources are provided in the Supplementary Material. It is important to note that the dataset includes variables of different types; most environmental variables are continuous, whereas species-related variables tend to be discrete. To accommodate the mixed data types, in tPC, PC and RestPC we use a conditional independence test based on the Bayesian Gaussian copula model as introduced in [22], using the Markov chain Monte Carlo sampler proposed by [22]; see the Supplementary Material, for details. This approach allows us to use the posterior distribution of the correlation matrix obtained from the Gaussian copula model to run each causal discovery algorithm several times, thereby incorporating uncertainty in the estimation of the correlation matrix into the estimation of the SCG. We drew 50 posterior samples of the correlation matrix and applied both PC and RestPC to each sample, obtaining one SCG and one partial causal order per method and sample. Each inferred order was then compared with the temporal reference order obtained by applying tPC to the two-time-point data. We used two notions of non-contradiction. The first counts only explicit reversals as contradictions: if *X* precedes *Y* in the reference order, a contradiction occurs only when the inferred order places *Y* before *X*. The second is stricter: non-contradiction requires the inferred order to also place *X* before *Y*. We report the mean and variance of these non-contradiction scores across the 50 samples. The significance level *α* for the (conditional) independence tests is set to 0.05.

Both single-time-point methods showed limited agreement with the causal order obtained from the two-time-point method, as expected due to the loss of temporal information. In terms of causal-order consistency, the non-contradiction rate was 0.81 ± 0.01 for RestPC versus tPC and 0.99 ± 0.01 for PC versus tPC. At first sight, this could suggest that PC performs better. However, this result is misleading, as it mainly reflects the fact that PC collapses most variables into the same level of the partial causal order. When considering the stricter agreement criterion, the rate drops to 0.14 ± 0.001 for RestPC and 0.01 ± 0.001 for PC. Thus, RestPC is more consistent with the reference order, although the overall level of agreement remains limited.

To discuss the results from an ecological perspective, we derived two consensus graphs: one from the 50 graphs obtained by PC and one from those obtained by RestPC. In each case, we retained only edges appearing in more than 50% of the samples, and then deduced the corresponding consensus partial causal order. Sensitivity analyses reported in the Supplementary Material show that the results are robust to the choice of the edge-retention threshold.

We pre-specified a small set of directional relations that are largely uncontroversial on ecological grounds independently of any causal discovery output: climate and non-human made land-cover variables are background drivers of bird abundance and therefore precede it, and altitude precedes climate. Among these 182 pre-specified relations, the RestPC consensus order satisfies 15 and contradicts 27 and has unresolved 140, while the PC consensus order satisfies 31 and contradicts 5 and has unresolved 146. Although informative, this pairwise evaluation is influenced by the relative sizes of the ecological groups. In particular, because the bird-abundance group contains substantially more variables than the environmental groups, a single environmental variable incorrectly placed downstream of the bird group generates many violated pairwise relations. Conversely, an incorrectly placed bird variable generates comparatively few violations.

To complement this analysis, we also examined the variables explicitly separated from the largest inferred causal-order level. Both PC and RestPC produced two causal-order levels: one large level containing most variables and one smaller level. PC separated two variables from the largest level, one in agreement with the prespecified ecological hierarchy and one in disagreement. In contrast, RestPC separated 4 variables, 3 in agreement with the hierarchy and one (`proportion of human-made land cover`) in disagreement. One plausible explanation for this disagreement is that land-cover variables are compositional, meaning their proportions are constrained to sum to a constant. This could mean that even after removing collinear cover classes, the remaining variables can retain a non-causal residual dependence, which is an instance of the faithfulness violations discussed in Section 2. Here, canceling or constraint-induced pathways can distort conditional independence relations. However, we did not formally test this hypothesis (e.g. via an isometric log-ratio transform of the composition). A further possibility is exact temporal determinism, whereby some variables remain unchanged across consecutive time points. An alternative, more mundane explanation is that the level-extraction procedure used to convert a partially resolved consensus order into discrete levels may place weakly identified variables at the end by default, independently of any genuine causal signal. We flag these possibilities without adjudicating between them and will leave a dedicated empirical test for future work.

Overall, neither RestPC nor PC consistently dominates the other; which method appears preferable depends on the criterion used for comparison. However, we emphasize that these comparisons reflect relative interpretability given our assumptions and pre-specified checks, not independent confirmation that PC and RestPC recovered the correct structure.

## 6 Structural relations among environmental, soil, and geochemical variables

In this section, we illustrate PC and RestPC on data related to terrestrial ecosystem components. We use environmental, soil physical, and soil chemical data from [17], derived from the soil campaign of the Land Use/Land Cover Area Frame Survey (LUCAS) [58]. The analysis includes 24, 930 distinct sampling locations (see details in the Supplementary Material). The selected variables capture different dimensions of the ecosystem, including environmental variables such as precipitation and temperature (from CHELSA database [14]); terrain characteristics such as elevation, slope, aspect, and distance to water (from Copernicus Digital Elevation Model); and geological characteristics (from the European Soil Database). In addition, we incorporate soil physical variables, soil chemical properties, and heavy metal concentrations (from the LUCAS campaign dataset). Because repeated measurements over consecutive years are not available for each location, we consider one sample per location, taking the last available year for each site corresponding to a single-time-point setting. See more details in the Supplementary Material.

We use the same conditional independence tests, the same significance level, and edge-retention procedure (sensitivity analyses in the Supplementary Material show robustness to the edge-retention threshold) as in the previous section. However, for this dataset, unlike in the previous section, only single time-point observations are available. Therefore we cannot use tPC on two-time-point data as a reference partial order. The comparison between PC and RestPC must instead be interpreted only in terms of ecological plausibility.

As in the previous section, we pre-specified a small set of directional relations that are plausible on physical or ecological grounds alone: terrain and parent-material variables (e.g., `>elevation`, `>slope`, `>aspect`) precede soil physical properties, which in turn precede soil geochemical and heavy-metal concentrations; and climate variables precede soil chemical properties. Among these 307 relations, the RestPC consensus order satisfies 159 and contradicts 91 and has 57 unresolved, while the PC consensus order satisfies 38 and contradicts 49 and has 220 unresolved. As above, this is a coarse plausibility check, not an independent validation of either method. More precisely, the PC order was weakly resolved: most variables were placed in one large intermediate class, providing limited insight into the ordering of terrain, climate, soil, and geochemical processes. In particular, only 1*/*9 terrain and parent-material variables appeared in the first level, namely `>distance to coast`. Only 1*/*11 downstream geochemical variables, `>manganese`, appeared in the final level. By contrast, RestPC produced a more ecologically interpretable order. It placed 5*/*9 background terrain and parent material, variables in the first level: `>aspect`, `>distance to coast`, `>distance to freshwater`, `>erosion`, and `>terrain ruggedness index`. It also placed 6*/*11 geochemical variables in the final level, namely `>arsenic`, `>cobalt`, `>manganese`, `>nickel`, `>antimony`, and `>zinc`.

Overall, RestPC better separates part of the upstream landscape and soil-context structure from a downstream geochemical block. However, the order is not fully mechanistically interpretable. In particular, the early positions of `>cadmium`, `>mercury`, and `>copper`, and the late position of `>snow-water equivalent`, remain difficult to explain ecologically.

## 7 Discussion

In this paper, we showed when and how constraint-based causal discovery can still be informative for single-time-point observational data generated by ecological dynamical systems. Although causal sufficiency over the DAG is violated in this setting because all processes are first-order self-dependent, the PC algorithm remains able to identify specific structures, namely *clustered-super-unshielded-colliders* (CSUCs). While CSUCs do not reflect exact variable-level causal relations, they provide partial process-level ordering information, indicating that some processes are down-stream of others. This is most informative when clusters are small or ecologically coherent; for large clusters, the statement may be too coarse for direct ecological interpretation. Nevertheless, other structural patterns are generally not identifiable from single-time-point data, and the FCI algorithm does not provide more informative results despite being designed to handle unmeasured confounding. We also introduced RestPC, a more efficient variant of PC tailored to the detect CSUCs. We supported these theoretical results with simulations studies.

In addition, we illustrated PC and RestPC on two real-world datasets. Some of the recovered relations were consistent with ecological expectations, but others were not fully mechanistically interpretable. Such deviations may arise for several reasons. First, ecological systems are affected by many unmeasured or imperfectly measured drivers, which may violate causal sufficiency over the SCG. Second, faithfulness may be difficult to justify in ecology, where multiple interacting pathways can plausibly cancel one another, making conditional independencies difficult to interpret; addressing such violations remains an important direction for future work [49, 16]. Third, consistency through time may fail, in which case single-time-point data become less informative about process-level relations. Nevertheless, this assumption remains weaker than the stationarity-type assumptions often used in causal discovery for time series, which additionally require causal strengths to remain unchanged over time. Fourth, although first-order self-dependence is usually ecologically plausible (especially when data are collected yearly [13]), violations of this assumption can weaken the conclusions; the implications of such violations are discussed in the Supplementary Material. Another notable practical concern is the nature of the variables. In real-world datasets, not all variables correspond to dynamical processes; for example, altitude in the STOC dataset and elevation in the LUCAS dataset are better viewed as static or slowly varying environmental variables. Such variables may make PC more informative than RestPC, as our sensitivity analyses demonstrate (see the Supplementary Material), since PC can exploit conditional independences beyond the dynamical-process interpretation. Finally, one specific limitation of the LUCAS dataset is spatial autocorrelation arising from the sampling design, which may violate the iid assumption underlying the conditional independence tests.

Most of the limitations discussed above are largely tied to assumptions that are difficult to test from observational data alone. Their plausibility should therefore be assessed with ecological expertise and, when possible, supported by substantive knowledge about the system under study. One way to improve interpretability is to incorporate background knowledge. The algorithms considered here allow such knowledge to be included directly, although we did not use it in this study in order to evaluate the methods on their own. Reliable ecological knowledge can help rule out implausible structures and guide the discovery process toward more realistic graphs, as is common in causal discovery in both non-dynamical settings [86, 52] and time-series settings [39, 51]. Similar strategies could be adapted to the single-time-point setting considered here.

Future work should investigate if process-level information beyond CSUCs can be recovered from conditional independence information at a single-time-point. It would also be interesting to extend this analysis to settings with partial temporal information, mixtures of static and dynamic variables, or alternative assumptions on the underlying dynamical system.

## Acknowledgments

We would like to thank our missing friend, Marc Ohlmann, for the early discussions we had with him about this work, as well as for his thorough proofreading of an early version of this work and for providing valuable feedback. We thank Pierre Gaüzère for providing the pre-processed real-world dataset from the French Breeding Bird Survey. We thank Simon Ferreira for his collaboration with D.B. on a project presented at the Tools for Causality Colloquium where they jointly applied the temporal PC algorithm to a two-points-in-time data version of the real-world dataset from the French Breeding Bird Survey, which included only environmental variables. We thank anonymous reviewers and guest editors for their helpful comments that substantially improved this paper. This work has been partially supported by the Horizon Europe Obsgession project (No: 101134954), the French National Research Agency under the France 2030 programme [Grant Number ANR-25-APDY-0001, DYNABIOD], the FRB-CESAB through the IMPACTS working group, and by the CIPHOD project (ANR-23-CPJ1-0212-01).

## Supplementary Material

### A FCI algorithm

In the main paper, we considered the FCI algorithm because it is a natural counterpart to PC in settings where unmeasured confounding may be present. This is particularly relevant for single-time-point data generated by dynamical systems, where unobserved past states can act as latent common causes of the observed variables. However, as shown in Theorem 1, in the setting considered here, FCI does not provide additional recoverable process-level information beyond the CSUCs already detected by PC. Since the main paper did not include a description of FCI, we provide one here for completeness.

The initial step of the FCI algorithm mirrors the behavior of the PC algorithm in deducing a skeleton. However, the skeleton inferred by this step may not necessarily correspond to the true skeleton, as the presence of hidden common causes could indicate that an edge between *X* and *Y* cannot be eliminated solely by conditioning on the neighbors of *X* and *Y*.

Within a MAG, a vertex *Z* is considered to be in the set of D-Sep(*X, Y*) if and only if *X* ≠ *Z* and there exists an undirected path linking *X* and *Z* such that every vertex along the path, excluding the endpoints, takes the form of a collider and is an ancestor of either *X* or *Y* [86]. The FCI algorithm employs an extended version of D-Sep, known as Possible-D-Sep(*X, Y*).

#### Definition 2

(Possible-D-Sep). *Z* ∈ *Possible-D-Sep*(*X, Y*) *if and only if there exists an undirected path π connecting X and Z in such a way that every subpath < A, B, C* > *along the path either constitutes a collider or forms a triangle*.

To identify those Possible-D-Sep relationships, FCI initiates the process by applying FCI-Rule 0, a counterpart to PC Step 2.1. Unlike PC Step 2.1, FCI-Rule 0 is capable of orienting bi-directed edges.

**FCI-Rule 0**. *Given an unshielded triple* 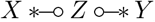, *if Z* ∉ *Sepset*(*X, Y*) *then* 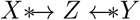. *Here, the asterisk (*\**) functions as a meta-symbol, serving as a wildcard to represent any of the three marks*.

After using FCI-Rule 0, FCI proceeds to eliminate edges using the Possible-D-Sep approach. It subsequently reorients all edges pocessing bi-directed loops and reapplies FCI-Rule 0. Subsequently, similar to the procedure in the PC algorithm, FCI continues to orient more edges by employing additional orientation rules. Further details about these rules can be found in [99].

### B Pseudocode of RestPC

#### Algorithm 1

`RestPC`.

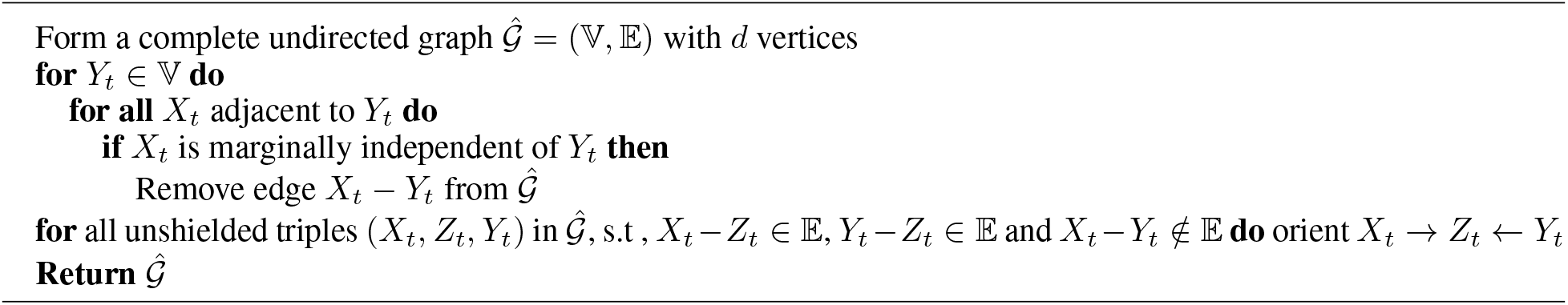

### C Other examples

Here we give two more examples (Figure 2) of time series with first-order self dependence. In the first example, we can see that both PC and FCI are able to find the cluster super unshielded colliders. In the second example, which is similar to the one given by [86], but here all the past is considered as hidden common cause, both PC and FCI give a fully connected undirected graph.

### D Proofs

#### Lemma 1

*Consider a SCG G*^*s*^, *and a compatible full-time DAG*. *If* ***consistency throughout time*** *(Assumption 2*.*3) and* ***first-order self dependence assumption*** *(Assumption 2*.*4) are satisfied and there exists an active path between X and Y in G*^*s*^, *then*

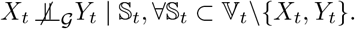

*Proof*. Let *π*^*s*^ = ⟨*V* ^1^ = *X, V* ^2^, …, *V*^*n*^ = *Y*⟩ be an active path between *X* and *Y* in the SCG *G*^*s*^. Since the path is active without conditioning, *π*^*s*^ contains no collider. Hence *π*^*s*^ is a trek, that is, it is the union of two possibly trivial directed paths with a common source [88]. Therefore, there exists *m* ∈ {1, …, *n*} such that

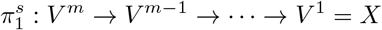

and

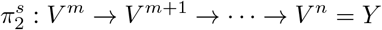

are directed paths in *G*^*s*^.

In what follows, we first construct a directed path in the full-time graph that is compatible with 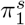, and then carry out the analogous construction for 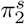. We subsequently show how the resulting paths can be combined to obtain an active path in the full-time graph. Finally, we use this construction to establish the main claim of the lemma. We recall that *t* is assumed to be sufficiently far from the lower boundary of the time domain so that all temporal indices introduced in the construction below are admissible.

#### Step 1: Lifting the branch from *V* ^*m*^ **to** *X*

For every edge

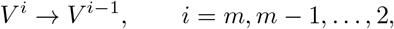

in 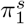, compatibility of *G*^*s*^ with the full-time DAG G implies that there exist a lag 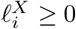 and an admissible time point *u* such that

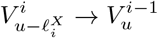

is an edge of G.

By consistency throughout time, this edge relation, with the same direction and lag, is present at every admissible time point.

Choose *r < t* such that all temporal indices constructed below are admissible. Set 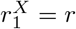 and define recursively

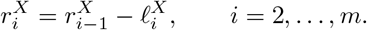

Assumption consistency throughout time then implies that

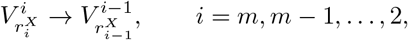

are edges of G. Hence

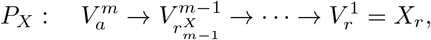

where 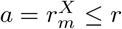, is a directed path in *G*.

If *m* = 1, the branch in 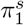 is trivial, and we simply take *P*_*X*_ to consist of the single vertex 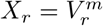, so that *a* = *r*.

#### Step 2: Lifting the branch from *V* ^*m*^ **to** *Y*

For every edge

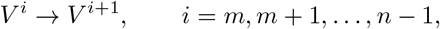

in 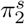, choose a corresponding lag 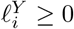. Set 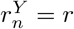 and define recursively, from *i* = *n* − 1 down to *m*,

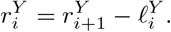

Again using consistency throughout time, we obtain a directed path

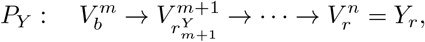

where 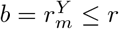.

If *m* = *n*, this branch is trivial, and we take *P*_*Y*_ to consist of the single vertex 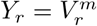, so that *b* = *r*.

#### Step 3: Giving the two lifted branches a common source

The two lifted copies of the SCG source *V* ^*m*^ need not occur at the same time. Let

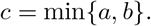

By first-order self dependence, for every process *U* and every admissible time *q*, the edge *U*_*q*−1_ → *U*_*q*_ belongs to *G*. In particular, G contains the directed self-dependence paths

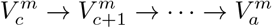

and

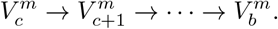

Concatenating these self-dependence paths with *P*_*X*_ and *P*_*Y*_, respectively, gives directed paths

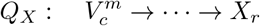

and

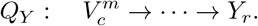

#### Step 4: Extending the endpoints to time *t*

First-order self dependence also gives

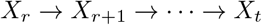

and

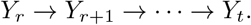

Appending these self-dependence paths to *Q*_*X*_ and *Q*_*Y*_ yields two directed paths

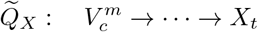

and

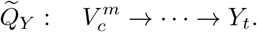

The two paths may have common vertices other than 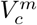, so their union is not necessarily a simple path. We therefore extract a simple path from them.

Let

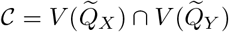

be their set of common vertices. This set is nonempty because 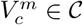.

Choose *Z* ∈ C to be the common vertex occurring last along 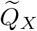, i.e., closest to *X*_*t*_ on that directed path.

We claim that no common vertex occurs strictly after *Z* on 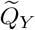. Suppose, for contradiction, that a common vertex *Z*^*′*^ occurs strictly after *Z* on 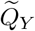. Since *Z*^*′*^ also lies on 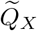, either:

1. *Z*^*′*^ occurs after *Z* on 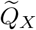, contradicting the choice of *Z*; or
2. *Z*^*′*^ occurs before *Z* on 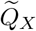.

In the second case, 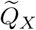 contains a directed path *Z*^*′*^ → · · · → *Z*, whereas 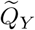 contains a directed path *Z* → · · · → *Z*^*′*^. Their union contains a directed cycle, contradicting the assumption that *G* is a DAG.

Thus, the subpaths

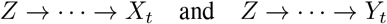

have no common vertex other than *Z*. Reversing the first subpath and concatenating it with the second gives the simple path

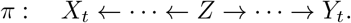

#### Step 5: Activity of the constructed path

The path *π* contains no collider. Every internal vertex on either directed branch has one incident arrowhead and one incident tail along the path, while both edges adjacent to *Z* have tails at *Z*. Therefore, *π* is active without conditioning.

Furthermore, every internal vertex of *π* occurs strictly before time *t*. Indeed, every vertex on the lifted SCG branches occurs at a time no later than *r < t*, and the internal vertices on the self-dependence paths from *X*_*r*_ to *X*_*t*_ and from *Y*_*r*_ to *Y*_*t*_ occur at times

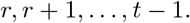

Thus, the only vertices of *π* occurring at time *t* are the endpoints *X*_*t*_ and *Y*_*t*_.

Let

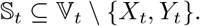

Every vertex in S_*t*_ occurs at time *t*, whereas every internal vertex of *π* occurs strictly before *t*. Therefore, no internal vertex of *π* belongs to S_*t*_.

Since *π* contains no collider and none of its internal non-colliders is conditioned upon, *π* is active given S_*t*_. Consequently,

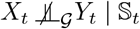

for every

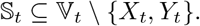

##### Theorem 2.

*Consider a distribution satisfying Assumptions 2*.*1, 2*.*2, 2*.*3, 2*.*4, and 2*.*5 and suppose we are given perfect conditional independence information about all pairs of variables* (*X*_*t*_, *Y*_*t*_) *in*V_*t*_. *Then the PC and FCI algorithms recover the components of the SCG and the same process-level orientation information. In particular, both algorithms recover the process-level orientation information represented by the collection of CSUCs*.

*Proof*. Since we assume the causal Markov condition and faithfulness, d-separation is equivalent to statistical conditional independence. So in the following, we will use d-separation to refer to both d-separation and conditional independence.

First, we prove that the PC algorithm is able to find graph components. If two vertices *X* and *Y* in the SCG belong to different components, then there is no active path between them, so *X*_*t*_ ⫫ _*G*_*Y*_*t*_ | ∅ given the Markov condition, faithfulness and causal sufficiency. Therefore, the PC algorithm would be able to separate *X*_*t*_ and *Y*_*t*_ in the first step of the algorithm, i.e., without conditioning on any vertex and so in the deduced SCG *X* and *Y* won’t be adjacent. Thus, the PC algorithm would find the different components in the SCG.

In the following we consider that the SCG contain one component. We assume that all the considered time series are first-order self dependent. In the first part, we consider skeleton construction. PC algorithm starts with the complete non oriented graph. We consider we have a pair of vertices (*X, Y*) in the SCG. We prove that PC removes *X* − *Y* if only if *X*_*t*_ ⫫_*G*_*Y*_*t*_ | ∅.

In the first step, the PC algorithm removes the edges between the vertices that are d-separated using an empty set. If process *X* and process *Y* are not related in the SCG neither directly nor indirectly (via common ancestors or intermediate causes), i.e., there is no active path between them, then ∀*t*_*i*_, *t*_*j*_, 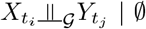, which means that *X*_*t*_ ⫫_*G*_*Y*_*t*_ | ∅. Thus, the PC algorithm will remove the edge between *X* and *Y* in the inferred SCG.

If process *X* and process *Y* are related directly or indirectly, i.e., there is an active path between them in the SCG then using Lemma 1 we obtain that 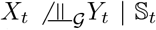, ∀S_*t*_ ⊂V_*t*_\{*X*_*t*_, *Y*_*t*_}. Hence, the PC algorithm would not be able to remove the edge between *X* and *Y* in the inferred SCG.

The second part concerns orientation. In the following, let us denote by Sep(*A, B*), the conditioning set that rendered *A* and *B* independent in the PC procedure. Let (X, ℤ, Y) be a CSUC. Consider arbitrary vertices

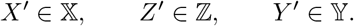

By condition (ii) of the definition of CSUCs, there exists an active path between *X*^*′*^ and *Z*^*′*^ and an active path between *Y* ^*′*^ and *Z*^*′*^. It follows from Lemma 1 that

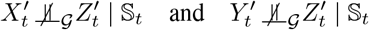

for every admissible instantaneous conditioning set S_*t*_. Consequently, PC does not remove the edges *X*^*′*^ − *Z*^*′*^ and *Y* ^*′*^ − *Z*^*′*^ during the skeleton phase. On the other hand, condition (i) implies that there is no active path between *X*^*′*^ and *Y* ^*′*^ in the SCG. By the causal Markov condition and faithfulness, 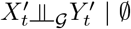. Therefore, PC removes the edge *X*^*′*^ − *Y* ^*′*^ using the empty separating set. Hence *X*^*′*^ − *Z*^*′*^ − *Y* ^*′*^ is an unshielded triple in the skeleton inferred by PC. Moreover, Sep(*X*^*′*^, *Y* ^*′*^) = ∅, and thus *Z*^*′*^ ∉ Sep(*X*^*′*^, *Y* ^*′*^). Step 2.1 of PC therefore orients *X*^*′*^ → *Z*^*′*^ ← *Y* ^*′*^. Since *X*^*′*^, *Z*^*′*^, and *Y* ^*′*^ were arbitrary, PC orients *X*^*′*^ → *Z*^*′*^ ← *Y* ^*′*^ for every

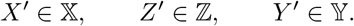

These triples are unshielded colliders in the graph inferred by PC, although they need not all be genuine unshielded colliders in the original SCG, because the inferred adjacencies may represent active paths rather than direct edges. Nevertheless, every CSUC contains at least one genuine unshielded collider in the SCG.

We now show that no subsequent PC orientation rule is applicable. For the first Meek rule, suppose that after Step 2.1 the inferred graph contained *A* → *B* − *C* with *A* not adjacent to *C*. Then *A B C* would be an unshielded triple. Since Sep(*A, C*) =, Step 2.1 would already have oriented *A* → *B* ←*C*, contradicting the assumption that *B* − *C* remains unoriented. Hence the first Meek rule cannot be applied. For each of the remaining Meek rules, its antecedent contains an unoriented edge together with directed edges produced at Step 2.1. Because every directed edge produced at Step 2.1 is an arrowhead into the middle vertex of an unshielded triple whose endpoints have empty separating set, the antecedent of any remaining rule would necessarily contain an unshielded triple that Step 2.1 should already have oriented at the previously unoriented edge within an unshielded triple. This contradicts the presence of that unoriented edge. Therefore, none of the subsequent Meek rules is applicable.

Since the FCI-algorithm start in the same way as the PC-algorithm, we can directly conclude that it can find different components and it will be able to prune the edges between the extremities of unshielded collider for which the extremities are marginally independent. After using the first phase of the PC algorithm, the FCI algorithm proceeds to pruning additional edges using Possible-D-sep sets. However, similarly to the argument given for the skeleton phase of the PC algorithm conditioning of Possible-D-sep set at time would not be yield any independence due to first-order self dependence.

We now consider the orientation phase. FCI initially represents every retained adjacency as *X* ∘−∘*Y*. Rule (R0) orients an unshielded triple as *X*, → *Z ←, Y* whenever *Z* ∉ Sep(*X, Y*). In the present setting, every missing adjacency is detected using the empty separating set. Thus, for every unshielded triple in the recovered skeleton, Sep(*X, Y*) =∅, and therefore *Z ∉* Sep(*X, Y*). It follows that (R0) orients every unshielded triple as a collider. Since PC and FCI recover the same skeleton and use the same empty separating sets, the arrowheads introduced by FCI Rule (R0) coincide with those introduced by Step 2.1 of PC. We now show directly that none of the subsequent FCI orientation rules unrelated to selection bias is applicable. The remainder of the proof assumes familiarity with the concepts and notation associated with the FCI algorithm introduced in [99]. After Rule *R*0, every unshielded triple *A* − *B* − *C* in the recovered skeleton is oriented as *A\** → *B* ←*\* C*, because *A* and *C* are nonadjacent and Sep(*A, C*) = ∅. Moreover, *R*0 introduces only arrowheads at the middle vertices of unshielded triples; it introduces no tail marks. Hence, immediately after *R*0, every endpoint mark is either a circle or an arrowhead. Suppose, for contradiction, that one of the subsequent rules *R*1, *R*2, *R*3, *R*4, *R*8, *R*9, *R*10 is applicable. Consider the first application of any of these rules. Before this first application, the graph contains only the marks produced by *R*0. For *R*1, suppose that the graph contains *A\** → *B* ∘−*\* C* such that *A* and *C* are not adjacent. Then *A* − *B* − *C* is an unshielded triple. Since Sep(*A, C*) = ∅, Rule *R*0 must already have oriented an arrowhead at *B* on the edge *B* − *C*, yielding *A \** →*B* ←*\* C*. This contradicts the circle mark at *B* required by the antecedent of *R*1. Therefore, *R*1 cannot be the first applicable rule. Rule *R*2 requires one of the configurations *A* → *B \** → *C* or *A \** → *B* → *C*. Each configuration contains at least one tail mark. However, before the first application of a rule subsequent to *R*0, no tail mark exists. Hence *R*2 cannot be the first applicable rule. For *R*3, suppose that its antecedent occurs. In particular, there exist vertices *A, B, C, D* such that *A \** → *B* ←*\*C, A \**−∘*D* ∘−*\* C, A* and *C* are not adjacent, together with the required adjacency between *D* and *B*. Since *A* and *C* are nonadjacent, *A* − *D* − *C* is an unshielded triple. Rule *R*0 must therefore already have oriented it as *A \** → *D* ←*\*C*. In particular, the endpoints at *D* on the edges *A* − *D* and *D* − *C* must be arrowheads, contradicting the circle endpoints required by the antecedent of *R*3. Thus *R*3 cannot be the first applicable rule. Rule *R*4 requires a discriminating path ⟨*θ*, …, *A, B, C*⟩ for *B*. By the definition of a discriminating path, every vertex between *θ* and *B* on this path is a parent of *C*. Consequently, the antecedent of *R*4 contains at least one directed edge into *C*, and therefore at least one tail mark. Since no tail marks are present before the first application of a rule subsequent to *R*0, no such discriminating path can occur. Hence *R*4 cannot be the first applicable rule. The antecedents of Rules *R*8, *R*9, and *R*10 also require directed or partially directed configurations containing at least one tail mark. For example, *R*8 requires a configuration containing a directed edge or a directed two-edge path, while *R*9 and *R*10 require a directed edge together with an uncovered potentially directed path. Since *R*0 introduces no tails, none of these antecedents can occur before some earlier rule has introduced a tail. But *R*1, *R*2, *R*3, and *R*4 have just been shown to be inapplicable. Therefore, *R*8, *R*9, and *R*10 cannot be the first applicable rule either. We have obtained a contradiction in every possible case. Hence none of the FCI orientation rules *R*1, *R*2, *R*3, *R*4, *R*8, *R*9, *R*10 is applicable after *R*0. Rules *R*5, *R*6, and *R*7, which concern tail and undirected-edge orientations associated with selection variables, are irrelevant under the assumption that selection bias is absent. Consequently, no additional endpoint marks are introduced after *R*0.

##### Corollary 2.

*Consider a distribution satisfying Assumptions 2*.*1, 2*.*2, 2*.*3, 2*.*4, and 2*.*5 and suppose we are given perfect marginal independence information about all pairs of variables* (*X*_*t*_, *Y*_*t*_) *in*V_*t*_. *Then the RestPC algorithm recovers the components of the SCG and the process-level orientation information represented by the collection of CSUCs*.

*Proof*. The proof follows from the proof of Theorem 1, where only the first two steps Step 1.1, Step 1.2 (pruning edges between vertices that are marginally independent) in the first phase of the PC algorithm and the Step 2.1 (orienting unshielded colliders) in the second phase were used. Which means, that by construction the RestPC algorithm would yield the same result as the PC algorithm.

### E Towards relaxation of the first-order self-dependence assumption

In this section, we investigate how the first-order self-dependence assumption can be relaxed, as well as settings in which the results of the paper no longer apply. We focus on PC and RestPC, although analogous results can be obtained for FCI.

The equivalence between RestPC and PC and the fact that they detect CSUCs relies on the fact that **an active path in the SCG can be represented by a compatible path in the full-time DAG whose internal vertices belong to the past**. The assumption that every time series causes itself at lag 1 guarantees this property, but it is stronger than necessary. We provide below a weaker condition based on the compatibility between self-causal lags and cross-series causal lags.

#### Proposition 1

(Past-only compatibility of active paths). *Let* G *be a stationary full-time DAG and let* G*s be its associated SCG. For each time series X, let L*_*X*_ ⊆ ℕ_>0_ *denote the non-empty set of self-causal lags of X, that is, l* ∈ *L*_*X*_ ⇐⇒ *X*_*t*−*l*_ → *X*_*t*_ *for all t. For every edge X* → *Y in G*^*s*^, *let*

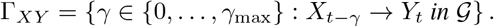

*Assume that there exists a common lag d* ℕ_>0_ *such that d* ∈ *L*_*X*_ *for every time series X, and that, for every edge X* → *Y in G*^*s*^, *there exists at least one γ* ∈ Γ_*XY*_ *such that γ* = *kd for some integer k* ≥ 0. *Then every active path in G*^*s*^ *admits a compatible active path in G between the time t instances of its endpoints, such that all non-endpoint vertices on the compatible active path occur at times strictly smaller than t*.

*Proof*. Let *d* be the common self-causal lag. Since *d* ∈ *L*_*X*_ for every time series *X*, stationarity implies *X*_*t*−(*m*+1)*d*_ → *X*_*t*−*md*_ for every *X* and every integer *m* ≥ 0. Therefore, each time series can be followed through the time points

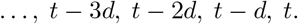

Now consider an edge *X* → *Y* in *G*^*s*^. By assumption, there exists a corresponding causal relation with lag *γ* = *kd* for some integer *k* ≥ 0. Hence, *X*_*t*−*kd*_ → *Y*_*t*_. By stationarity, the same causal relation also exists at earlier times: *X*_*t*−(*m*+*k*)*d*_ → *Y*_*t*−*md*_ for every integer *m* ≥ 0.

Thus, every edge of *G*^*s*^ has at least one representation using only time points whose differences are multiples of *d*. Moreover, between two consecutive such time points, every time series causes itself: *X*_*t*−(*m*+1)*d*_ → *X*_*t*−*md*_.

Consider now only these time points and, for each edge of *G*^*s*^, one corresponding causal relation whose lag is a multiple of *d*. The resulting full-time DAG has the same SCG *G*^*s*^. If a time difference of *d* is regarded as one time step, every time series causes itself at lag 1 in this DAG.

The original argument for lag-1 self-causation can therefore be applied. Hence, every active path in *G*^*s*^ has a compatible active path between the time-*t* instances of its endpoints, with all non-endpoint vertices occurring strictly before *t*.

Since all causal relations used in this compatible active path also belong to *G*, the same active path exists in *G*. This proves the result.

□

#### Remark about the relation with the original lag-1 assumption

The original assumption is recovered as the particular case *d* = 1. Indeed, if every time series causes itself at lag 1, then 1 ∈ *L*_*X*_ for every *X*, and the first condition of Proposition 1 is satisfied with *d* = 1. In this case, the condition on the cross-series causal lags imposes no additional restriction. For any *γ* ∈ {0, …, *γ*_max_}, we can write *γ* = *γ ×* 1. Hence every possible causal lag is compatible with *d* = 1, regardless of the value of *γ*_max_. Therefore, the original lag-1 setting is fully contained in the setting of Proposition 1.

#### Remark about the strict relaxation of lag-1 self-causation

The new condition also includes models in which no time series causes itself at lag 1. For example, suppose that 2 ∈ *L*_*X*_ for every time series *X*, while 1∉ *L*_*X*_. The common temporal grid can then be defined by *d* = 2. The condition of Proposition 1 requires each edge *X* → *Y* of the SCG to admit at least one causal realization at a lag of the form *γ* = 2*k*, that is, at lag 0, 2, 4, 6, For instance, the relations

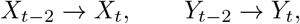

together with

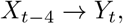

satisfy the assumptions even though neither *X* nor *Y* has a lag-1 self-causal relation. Thus, the proposed condition strictly extends the class of full-time causal graphs covered by the original lag-1 assumption.

#### Remark about the non restriction on all causal lags

Importantly, the condition does not require every causal lag in Γ_*XY*_ to be a multiple of *d*. It only requires that, for each edge *X* → *Y* in the SCG, at least one causal realization of that edge be compatible with the common temporal grid. Thus, Γ_*XY*_ may also contain causal relations at other lags. For example, with *d* = 2, a set such as Γ_*XY*_ = 1, 2, 5 satisfies the condition because it contains the compatible lag *γ* = 2.

#### When the sufficient condition is not satisfied

The conditions of Proposition 1 are sufficient, but not necessary, for RestPC and PC to recover the same graphical information. When they are satisfied, every active path in the SCG admits a compatible active path in the full-time DAG whose non-endpoint vertices occur strictly before the observation time. Under the remaining assumptions of the original result, RestPC and PC therefore recover the same connected components and CSUCs. Conversely, if the conditions of Proposition 1 are not satisfied, this does not imply that RestPC and PC necessarily return different results. Their behavior then depends on the temporal structure of the full-time DAG. In particular, systems without self-dependence and systems with self-dependence beyond lag 1 lead to different situations, which we discuss below. A summary of these situations is given in Table 2.

**Table 2:** Summary of the relation between the sufficient condition of Proposition 1, the outputs of PC and RestPC, and the corresponding recovery guarantees under different temporal settings.

| Setting | Condition of Prop. 1 | Relation between outputs | Guarantee / interpretation |
| --- | --- | --- | --- |
| Common self-causal lag $d$ and, for every SCG edge, at least one compatible lag $\gamma = kd$ | Satisfied | PC = RestPC | Both recover the connected components and CSUCs under the remaining assumptions of the original result. |
| No instantaneous cross-series causal relations and self-dependence lag is greater than cross-series lags | Not applicable | PC = RestPC | PC and RestPC remain equivalent. However, their common output need not recover the connected components and CSUCs of the SCG. |
| No instantaneous cross-series causal relations and no self-dependence | Not applicable | PC = RestPC | PC and RestPC remain equivalent. However, their common output need not recover the connected components and CSUCs of the SCG. |
| Only instantaneous cross-series causal relations and no self-dependence | Not applicable | PC may be more informative | The problem reduces to standard cross-sectional causal discovery. Under the usual assumptions, PC recovers the true CPDAG, whereas RestPC does not generally have the same guarantee. |
| Instantaneous and time-lagged causal relations and no self-dependence | Not applicable | PC may be more informative | PC can exploit additional conditional independences, but its output has not been characterized and cannot generally be interpreted as the CPDAG of the SCG. |
| Deterministic temporal dependence | Not covered by the proposition | PC may be more informative | When the past of a time series is deterministic, it does not introduce additional stochastic temporal dependence at the observation time. From the perspective of contemporaneous causal discovery, such variables can therefore behave similarly to variables in a cross-sectional i.i.d. setting. Consequently, the equivalence between PC and RestPC is no longer guaranteed: PC may more closely recover the cross-sectional ground truth, whereas RestPC may return a different graph. |

**Table 3:** Percentage of detecting the correct skeleton and F1-score on adjacencies and orientations using the PC algorithm and the RestPC algorithm.

|  |  | F1-a-true | F1-o-true | F1-a-expected | F1-o-expected |
| --- | --- | --- | --- | --- | --- |
| 10 variables | PC | $0.53 \pm 0.01$ | **$0.29 \pm 0.01$** | $0.62 \pm 0.01$ | $0.23 \pm 0.01$ |
| | FCI | **$0.54 \pm 0.01$** | $0.28 \pm 0.01$ | $0.57 \pm 0.01$ | $0.21 \pm 0.01$ |
| | RestPC | $0.41 \pm 0.01$ | $0.13 \pm 0.01$ | **$0.95 \pm 0.01$** | **$0.7 \pm 0.03$** |
| 20 variables | PC | **$0.28 \pm 0.01$** | **$0.23 \pm 0.01$** | $0.46 \pm 0.01$ | $0.1 \pm 0.01$ |
|  | FCI | — | — | — | — |
| | RestPC | $0.21 \pm 0.01$ | $0.05 \pm 0.01$ | **$0.93 \pm 0.01$** | **$0.59 \pm 0.01$** |

#### Self-dependence beyond lag 1 and not satisfying the condition of Proposition 1

We now consider systems in which the processes are self-dependent, but their self-dependence is not necessarily of first order. In this setting, the guarantees obtained under lag-1 self-dependence do not generally hold. First-order self-dependence allows the vertices of an active path in the SCG to be temporally synchronized so as to induce a compatible active path between instantaneous variables. When self-dependence occurs only at larger lags, such a synchronization may fail. For example, consider a stationary full-time DAG containing

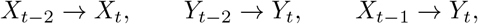

but no first-order self-dependence relation

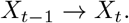

The corresponding SCG contains the edge *X* → *Y*, since a past value of *X* directly causes a future value of *Y*. Nevertheless, *X*_*t*_ and *Y*_*t*_ may be marginally independent. Indeed, *X*_*t*_ depends on past values of *X* whose time indices have the same parity as *t*, whereas the effect of *X* on *Y*_*t*_ is transmitted through *X*_*t*− 1_, which belongs to the opposite parity class. In the absence of any additional causal path connecting these two classes, there is no active path between *X*_*t*_ and *Y*_*t*_, and therefore

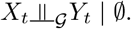

Consequently, RestPC removes the adjacency between *X* and *Y* during its marginal-independence step. PC also removes this adjacency during its skeleton-search phase, since the variables are already marginally independent. Thus, in this example, PC and RestPC still return the same result, but both fail to recover the process-level edge *X* → *Y*. More generally, when the self-causal and cross-series lags are not temporally compatible, instantaneous conditional-independence information may fail to reveal genuine process-level adjacencies. This may split connected components of the SCG and prevent the recovery of some CSUCs. Hence, the failure of the sufficient condition does not necessarily imply that PC and RestPC differ; it may instead imply that both algorithms miss the same process-level causal information.

#### Systems without self-dependence

When the considered dynamic system contains no self-dependence, three main situations can be distinguished. If all causal relations are instantaneous, the problem reduces to classical causal discovery from i.i.d. observations. Under Assumptions 2.1 and 2.2 and causal sufficiency, PC asymptotically recovers the true CPDAG. In this setting, RestPC does not in general recover the same information as PC, since its restricted use of conditional-independence tests is no longer justified by self-dependence. If all causal relations have strictly positive temporal lags, instantaneous observations alone do not in general identify the lagged causal structure represented by the SCG. In particular, an edge *X* → *Y* in the SCG need not induce an adjacency between *X*_*t*_ and *Y*_*t*_, while instantaneous variables may still be dependent because of common causes in the past. Consequently, both PC and RestPC may fail to recover genuine process-level relations from instantaneous observations alone. Finally, when instantaneous and time-lagged causal relations coexist, PC may recover more conditional-independence information than RestPC, since it also considers conditional independences beyond the marginal ones used by RestPC. However, the interpretation of the graph returned by PC in this setting is not currently characterized. In particular, because contemporaneous dependencies may be induced by both instantaneous and lagged causal relations, the output of PC cannot in general be interpreted as the CPDAG of the SCG. Thus, even if PC is more informative than RestPC, it remains unclear which features of the underlying dynamic causal structure are represented by its output.

#### Deterministic temporal dependenc

We finally consider mixed systems in which some processes remain self-dependent, whereas others exhibit deterministic temporal persistence. In this setting, observing only the variables at time *t* may simplify the problem faced by PC, while the equivalence between PC and RestPC is no longer guaranteed. For example, consider three processes *E, X*, and *Y*, where *X* and *Y* are genuinely self-dependent, while the environmental driver *E* satisfies

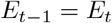

deterministically. Suppose further that the full-time causal structure contains

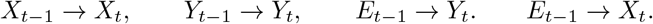

For instance, the structural equation of *Y*_*t*_ and *X*_*t*_ may be written as

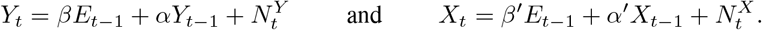

Since *E*_*t*−1_ = *E*_*t*_, this equation is observationally equivalent, at time *t*, to

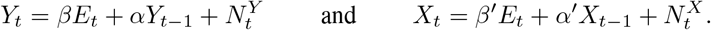

Thus, conditioning on *E*_*t*_ would render *X*_*t*_ and *Y*_*t*_ independent. Therefore, from the perspective of PC, which is applied only to variables at time *t*, the lagged contribution of *E* behaves as an instantaneous relation involving the observed variable *E*_*t*_. Consequently, part of the dynamic problem reduces to an ordinary cross-sectional causal-discovery problem, making the corresponding relations easier for PC to recover. This simplification does not provide the same guarantee for RestPC. Its restricted search is justified through the temporal self-dependence that is stochastic. In a mixed setting where some processes behave deterministically while others remain genuinely self-dependent, these conditions need not hold. Hence, PC and RestPC are no longer guaranteed to recover the same graphical information, and PC may benefit more directly from the cross-sectional structure induced by the deterministic variables.

Many of the situations discussed in this section will also be investigated empirically in Section F.1 using simulated data.

### F Extension of the simulation based experiments

This section provides additional experiments assessing the robustness, scalability, computational efficiency, and applicability of the proposed approach beyond the main experimental setting. We first investigate its robustness to deviations from the first-order self-dependence assumption, including settings covered by, as well as settings violating, the sufficient condition introduced in Proposition 1. We then evaluate the methods on larger causal graphs to assess how their performance evolves with increasing problem size. Next, we report and compare their computational times. Finally, we consider the nonlinear Lotka-Volterra dynamical system, providing an evaluation in a setting that departs from the linear data-generating mechanisms considered in the main experiments.

#### F.1 Robustness check against the first-order self-dependence assumption

In this section, we first conduct a sensitivity analysis comparing the first-order self-dependence setting with settings that violate both this assumption and the sufficient condition of Proposition 1. In particular, we fix the lag of every cross-series causal relation to 1 and vary the self-dependence lag of all time series between 1 and 3. Under this design, the condition of Proposition 1 is satisfied only when the self-dependence lag is 1. Indeed, for self-dependence lags 2 and 3, the cross-series lag *γ* = 1 is not a multiple of the common self-causal lag. For each setting, we perform 5000 simulation runs and report the average performance across runs. The results are reported in Figure 3. As expected, PC, FCI and RestPC achieve their best performance when the self-dependence lag is 1, corresponding to the setting covered by both the original first-order self-dependence assumption and Proposition 1. When the self-dependence lag is increased to 2 or 3, while keeping the cross-series lag fixed to 1, performance generally deteriorates substantially. The extent of this deterioration, however, depends on the considered causal structure and graphical feature. In particular, some expected adjacencies remain partially recoverable, whereas the recovery of the true adjacencies and orientation information is strongly degraded in most settings. Interestingly, PC, FCI and RestPC exhibit very similar behavior across all considered self-dependence lags. Thus, violating the sufficient condition of Proposition 1 does not necessarily imply that PC, FCI and RestPC return different results. In these simulations, both algorithms instead tend to lose the same process-level causal information.

We next consider settings that violate the first-order self-dependence assumption but still satisfy the sufficient condition of Proposition 1. We fix all cross-series causal relations to be instantaneous, i.e., *γ* = 0, and vary the common self-dependence lag between 1 and 3. Since 0 = 0 × *d* for any common self-causal lag *d*, all considered settings satisfy the condition of Proposition 1. As before, each setting is evaluated over 5000 simulation runs and the reported results correspond to averages across runs. The results are shown in Figure 4. In contrast to the previous experiment, performance remains remarkably stable as the self-dependence lag increases from 1 to 3. PC, FCI and RestPC obtain almost identical results across the different causal structures and graphical features, and no systematic deterioration is observed when moving away from first-order self-dependence. These results are consistent with Proposition 1 and illustrate that first-order self-dependence is not required for the equivalence between PC and RestPC when the temporal compatibility condition is satisfied.

**Figure 4.**
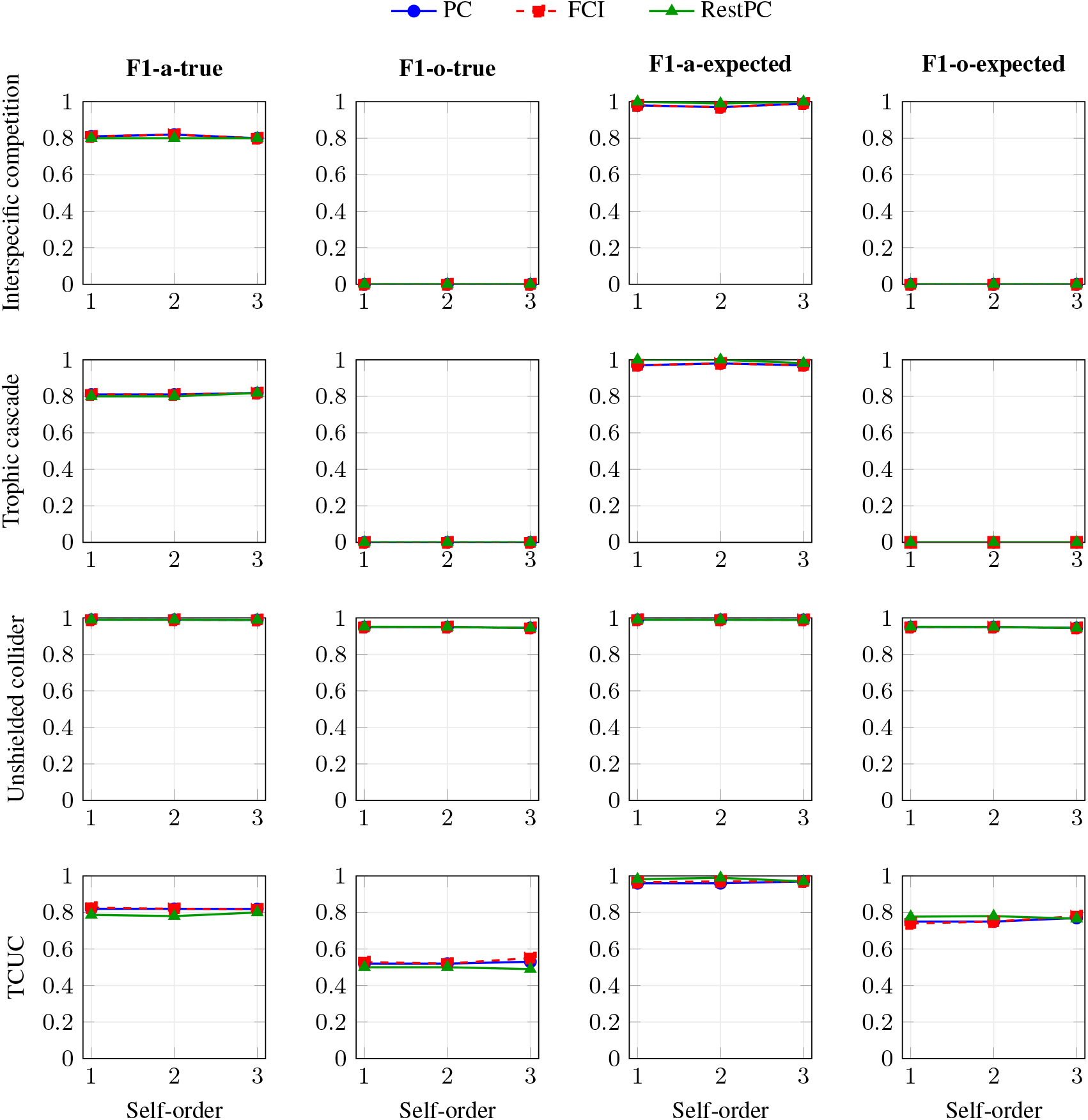
Sensitivity analysis to the self-dependence lag where TCUC is trophic cascade with unshielded collider. The lag of every cross-series causal relation is fixed to 0, while the self-dependence lag is varied from 1 to 3. All settings satisfy the condition of Proposition 1. Performance is reported for PC, FCI, and RestPC across the different causal structures and graphical features considered.

Finally, we consider settings that violate the condition of Proposition 1 by introducing deterministic temporal dependence for only a subset of variables. The results are reported in Figure 5. As expected, PC and FCI move closer to the ground truth as more variables are affected by deterministic temporal dependence. In these settings, RestPC no longer behaves similarly to PC and FCI, illustrating that once the sufficient condition is violated, the equivalence between RestPC and PC or FCI is no longer guaranteed.

**Figure 5.**
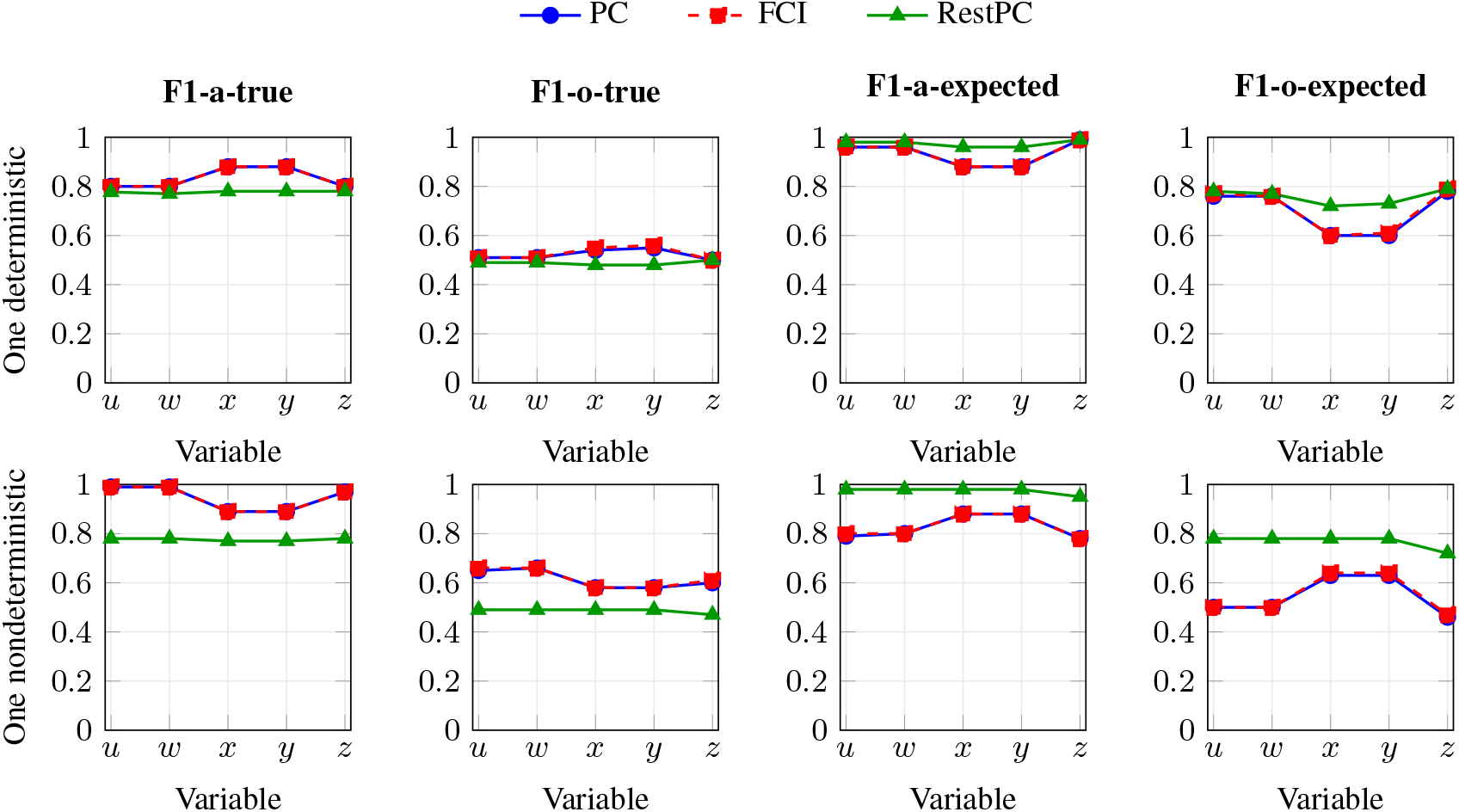
Sensitivity analysis to deterministic temporal dependence in the trophic cascade with unshielded collider setting. In the first row, one variable at a time is made deterministic, while all other variables retain their original data-generating mechanism. In the second row, the opposite setting is considered: all variables are made deterministic except one, with the horizontal axis indicating the variable that remains non-deterministic. Performance is reported for PC, FCI, and RestPC for each choice of variable.

#### F.2 Simulations based on larger graphs

In the main paper, we focused on settings with 3 or 5 variables. Here, we extend the analysis to larger settings with 10 and 20 variables. The simulation design is the same as before, except that, due to the increased computational cost, we ran 10 repetitions instead of 1000. The results show that, when performance is evaluated with respect to the true underlying graph, PC and FCI achieve slightly higher F1-scores than RestPC for both adjacencies and orientations. However, these scores remain relatively low and decrease as the number of variables increases, suggesting that recovering the full underlying structure from single-time-point data becomes increasingly difficult. In contrast, RestPC performs substantially better when evaluated against the expected graph: it achieves high F1-scores for adjacencies, equal to (0.95) for 10 variables and (0.93) for 20 variables, and also improves orientation recovery compared with PC. This is consistent with the objective of RestPC, which is not to recover the full micro-level causal graph, but rather the causal information that is identifiable at the process level. Finally, FCI could not be evaluated for 20 variables because its running time exceeded the predefined computational limit of 500 seconds per run.

#### F.3 Computation time

Figure 6 reports the average computation time of PC, FCI, and RestPC as the number of variables increases. Overall, RestPC remains substantially faster than both PC and FCI when increasing the number of variables. While the running time of PC increases markedly when moving to 20 variables, RestPC remains below one second on average. FCI also becomes computationally expensive as the number of variables increases; in particular, the result for 20 variables is not shown because the predefined computational time limit of 500 seconds per run was exceeded. These results suggest that RestPC provides a more scalable alternative in this setting, especially when the number of variables becomes larger.

**Figure 6.**
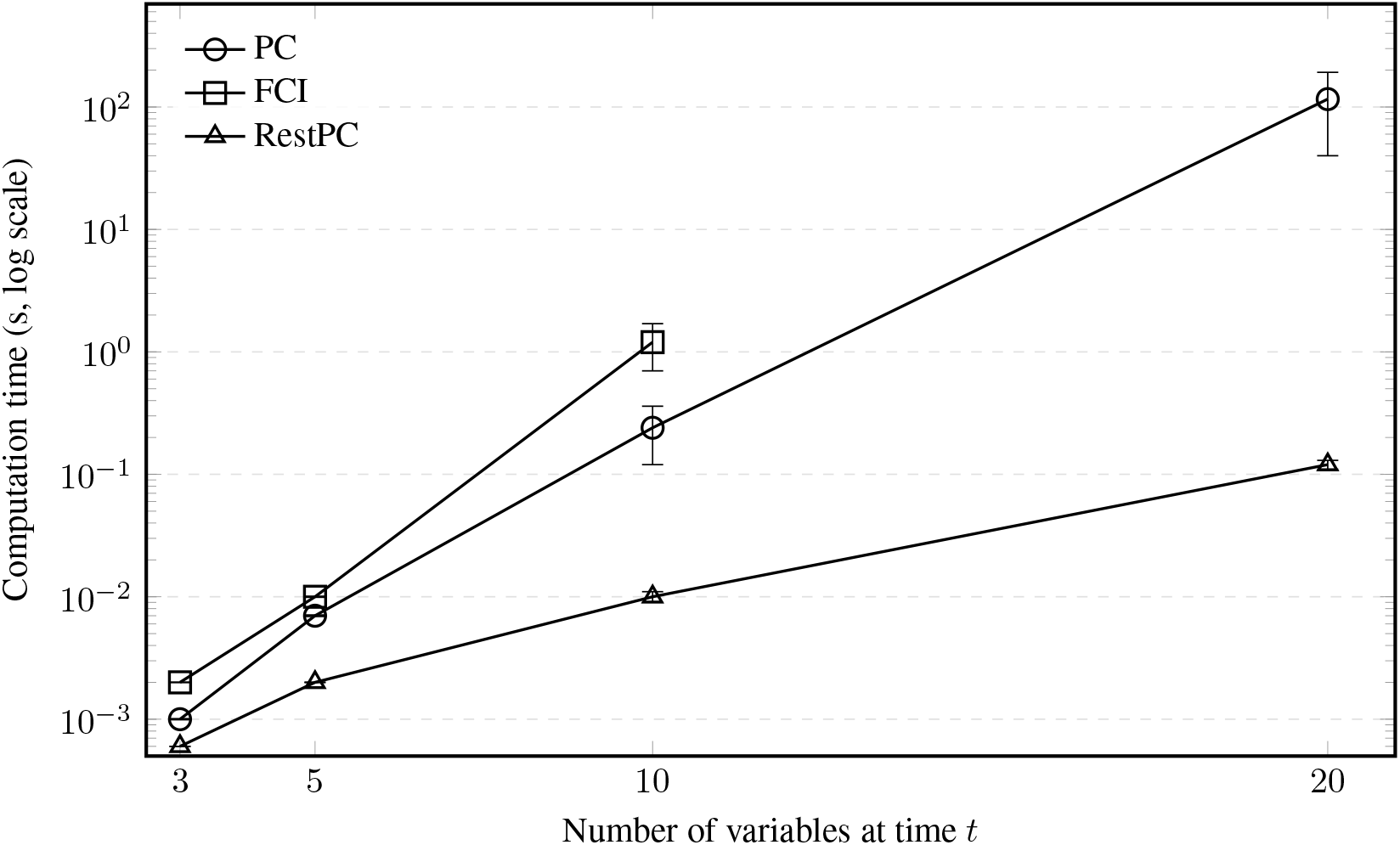
Average computation time of PC, FCI, and RestPC as a function of the number of variables. The y-axis is shown on a logarithmic scale. Error bars represent one standard deviation. The FCI result for 20 variables is not shown because it exceeded the predefined computational time limit.

#### F.4 Simulations based on Lotka Volterra model

We present results obtained from simulating data using a multi-species extension of the Ricker model introduced by [69]. This model, analogous to the generalized Lotka Volterra (GLV) with abiotic control, simplistically represents the primary processes within a complex ecological system. It incorporates feedback loops and self-loops, where species abundance depends on the previous time step’s abundance. Moreover, it accommodates non-linear species interdependencies.

We adopt this model as it is commonly used in ecological studies. The Ricker model with abiotic control in discrete time for the abundance of species *i* at time step *t* is given by:

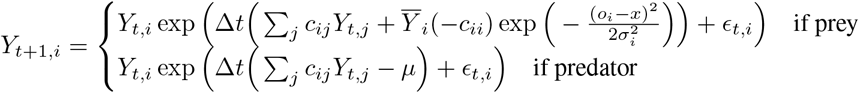

Here, *Y*_*t,i*_ represents the abundance of species *i* at time *t* and the upper equation is used when species *i* is prey, while the lower is used when species *i* is a predator. Notably, 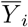 represents the abundance of species *j* in the stationary state, *c*_*ij*_ signifies the effect of species *i* on species *j, ϵ*_*t,i*_ is an independent and identically distributed (i.i.d) Gaussian random variable with variance *σ*_*r*_, *o*_*i*_ denotes the niche optimum for species *i, x* represents the environmental variable, and *µ* stands for the predator extinction rate.

We conducted simulations for 5 species, with fixed parameters: environmental variable *x* = 0.5, *T* = 1000 time steps, *µ* = 0.05, *σ*_*r*_ = 0.2, *o*_*i*_ randomly assigned between 0 and 1 for each species, and the interaction matrix derived from a randomly sampled graph *G* with 3 trophic levels and full connectivity. All links in the SCG are bi-directed (but the full-time graph is acyclic), and interaction strengths are randomly sampled for all interactions.

We applied PC, FCI, and RestPC algorithms to the data produced by this simulation, using a kernel-based conditional independence test [100] with a significance level of 0.05. We evaluated the methods by comparing the inferred graphs with both the true and expected SCGs. Since all edges are bi-directed (feedback loops), the expected SCG for this simulation is a fully connected undirected graph.

Table 4 displays the results of PC, FCI, and RestPC on the Lotka Volterra simulation. Observing the results, we find that RestPC excels at inferring the skeleton of the true SCG, outperforming FCI and PC. None of the methods, however, can precisely infer the true SCG. For orientations, as expected all methods perform poorly. Moreover, F1-a-e is better for all methods, with RestPC exhibiting the highest performance. Due to the absence of CSUCs, F1-o-e is not applicable, as there is only one component in the considered graphs.

**Table 4:** Percentage of detecting the correct skeleton and F1-score on adjacencies and orientations using the PC algorithm, the FCI algorithm and the RestPC algorithm on data simulated by the Lotka Voltera model. F1-o-expected is reported as “NA” for the interspecific competition and trophic cascade scenarios because, in these cases, the expected SCG contains no orientable structure.

|  |  | F1-a-true | F1-o-true | F1-a-expected | F1-o-expected |
| --- | --- | --- | --- | --- | --- |
| Lotka Volterra | PC | $0.12 \pm 0.03$ | $0.01 \pm 0.01$ | $0.29 \pm 0.03$ | NA |
| | FCI | $0.12 \pm 0.03$ | $0.09 \pm 0.03$ | $0.29 \pm 0.02$ | NA |
| | RestPC | $0.46 \pm 0.02$ | $0.09 \pm 0.03$ | $0.77 \pm 0.02$ | NA |

In conclusion, these results align with our theoretical framework. PC and FCI algorithms exhibit closer alignment with the expected SCG. The graph inferred by RestPC more closely resembles the expected graph than those of PC and FCI, as expected from the algorithm’s definition. Moreover, RestPC performs better compared to the true SCG, which can be attributed to its reduced need for conditional independence tests, minimizing opportunities for errors. This performance could be further improved by considering the trophic level background knowledge, which was not feasible due to the small scale of the simulation study.

### G Real-data analysis

Both datasets combine different types of data, so we used the Copula PC approach introduced by [22]. This approach uses a Bayesian Gaussian copula model to obtain samples from the posterior distribution of the correlation matrix.

Gaussian copula model:

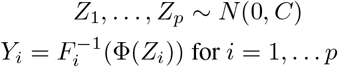

Gibbs sampling based on extended rank likelihood [22] with the following prior:

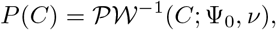

where *P* (Σ) = ^−1^(Σ, Ψ_0_, *ν*) and 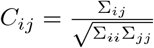. For each dataset, we ran two MCMC chains of 15, 000 iterations each, discarding the first 7, 500 iterations as burn-in for the first dataset and 5000 for the second dataset. Convergence was assessed using Gelman–Rubin diagnostics [32] and visual inspection of the trace plots.

For each setting (singe-time-point and two-time-point) we use the procedure suggested in [22]:

- *C*_1_, *C*_2_, … *C*_*m*_ ~ *P* (*C* | *Y*), where *m* = 50
- Compute effective sample size 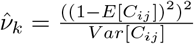 for all *Ci,j, i < j* and 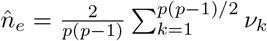
- Apply RestPC, PC or tPC on *C*_1_, *C*_2_, … *C*_*m*_ using 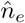 and significance level *α* = 0.05
- Compute *Ĝ*_*m*_ for each *m* and deduce corresponding SCG 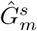.
  1. For tPC, PC and RestPC, computed the *Ĝ\** by keeping edges that appear with a probability higher than a threshold *β* (*β* = 0.5), deduce corresponding SCG *Ĝ*^*s\**^, then compute the partial causal order.
  2. For the two-time-point setting, for PC and RestPC, we compute the partial causal order for each 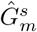. We then estimate the measure between the partial causal order obtained from 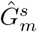 for each model and partial causal order in *Ĝ*^*s\**^ obtained from tPC.

In the following, for each dataset, we provide additional details on the data and preprocessing steps, report the partial order inferred by each algorithm, and conduct a sensitivity analysis with respect to the sparsity-threshold hyperparameters used by PC and RestPC. Finally, we perform a negative-control analysis following the approach of [67].

#### G.1 More detail and preprocessing of each dataset

##### STOC

All variables of this dataset, their definitions and sources are listed in Table 5. We used abundance data on breeding birds from the French Breeding Bird Survey (Suivi Temporel des Oiseaux Communs, STOC), a standardized monitoring program launched in 2001, in which skilled volunteer ornithologists identify breeding birds by song or visual contact each spring (see [54] for more details on the sampling methodology). We used the STOC bird monitoring dataset, together with the associated climatic variables, provided in [31]. We applied the same preprocessing steps, retaining 30 most abundant species that occurred in more than 50% of the sampled site-year combinations. Unlike the original study, however, we retained birds whose total abundance represented more than 0.5% of the total abundance across the community matrix (instead of 1%). Moreover, we did not fill missing observations, instead we retained the consecutive years with the most fully observed sites. Within these consecutive years we retained only fully observed sites. We analyze the interactions among 30 different bird species, the most abundant ones, incorporating 6 environmental variables including climate characteristics such as temperature, altitude and precipitation, and landcover characteristics such as habitat artificialization, water, agricultural areas, and forests. The STOC dataset spans at most 20 time-points, with each time-point representing a distinct year and collected from 1312 locations. Out of 1312 sites, we only included 824 sites with available consecutive data for the two years. The most recent time-point was used for single-time-point data, while the last two consecutive time-points were used for two-time-point data. For comparison, we also apply tPC to two-time-point data. We do not consider the resulting estimates as ground truth; instead, we use them to provide a complementary perspective on our findings from single-time-point data. We used land cover data from CORINE Land Cover (http://www.eea.europa.eu/data-and-maps/data/corine-land-cover-2006-raster) and climatic data extracted from CHELSA (https://chelsa-climate.org/, v.2.1).

**Table 5:**
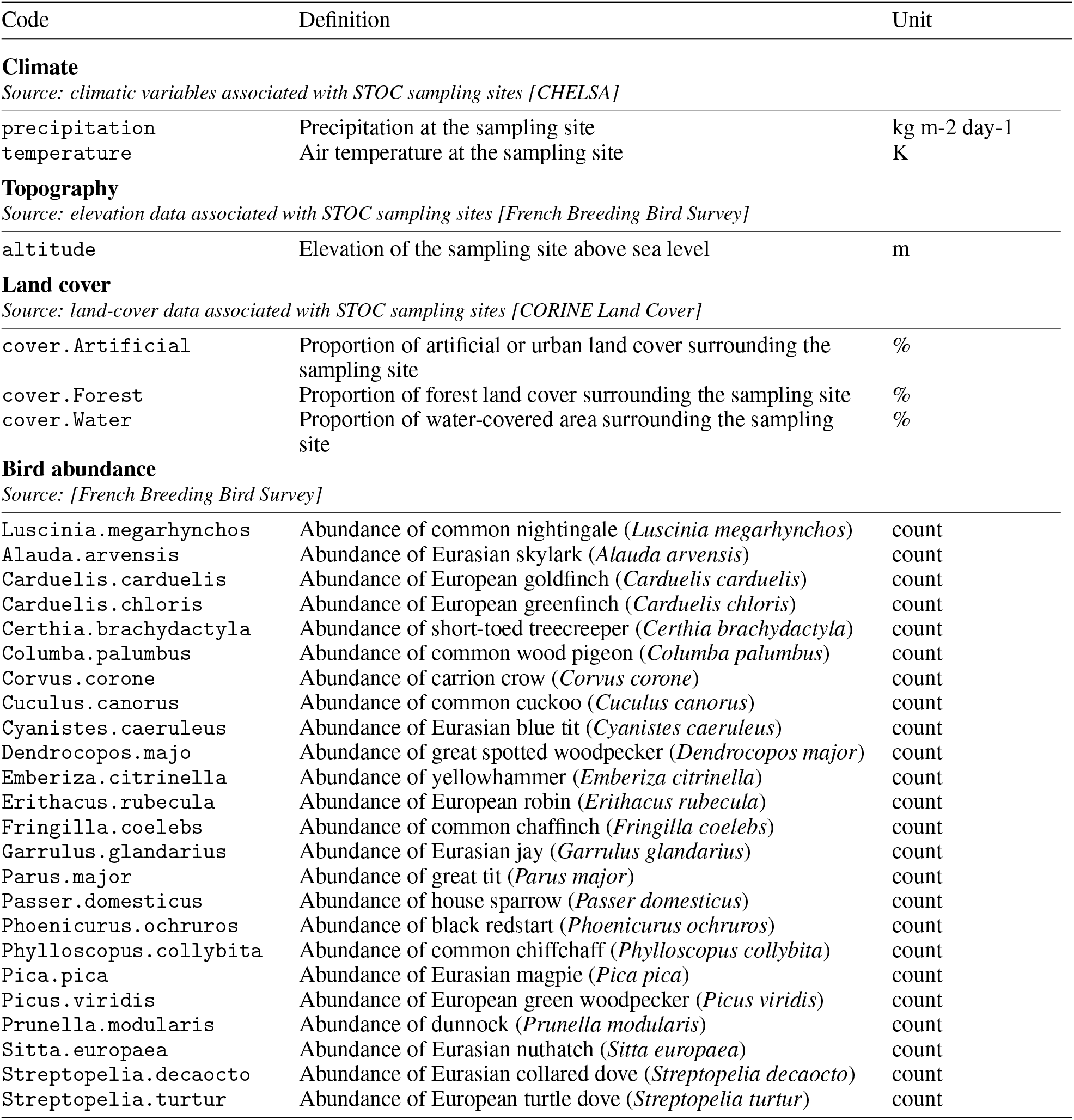
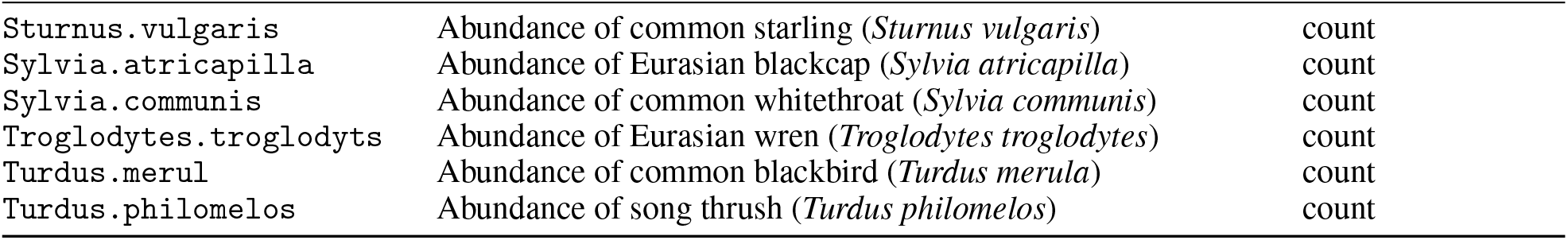
Variable definitions and data sources, grouped by feature type for the STOC dataset.

| Code | Definition | Unit |
| --- | --- | --- |
| **Climate** |  |  |
| *Source: climatic variables associated with STOC sampling sites [CHELSA]* |  |  |
| precipitation | Precipitation at the sampling site | kg m<sup>-2</sup> day<sup>-1</sup> |
| temperature | Air temperature at the sampling site | K |
| **Topography** |  |  |
| *Source: elevation data associated with STOC sampling sites [French Breeding Bird Survey]* |  |  |
| altitude | Elevation of the sampling site above sea level | m |
| **Land cover** |  |  |
| *Source: land-cover data associated with STOC sampling sites [CORINE Land Cover]* |  |  |
| cover.Artificial | Proportion of artificial or urban land cover surrounding the sampling site | % |
| cover.Forest | Proportion of forest land cover surrounding the sampling site | % |
| cover.Water | Proportion of water-covered area surrounding the sampling site | % |
| **Bird abundance** |  |  |
| *Source: [French Breeding Bird Survey]* |  |  |
| Luscinia.megarhynchos | Abundance of common nightingale (*Luscinia megarhynchos*) | count |
| Alauda.arvensis | Abundance of Eurasian skylark (*Alauda arvensis*) | count |
| Carduelis.carduelis | Abundance of European goldfinch (*Carduelis carduelis*) | count |
| Carduelis.chloris | Abundance of European greenfinch (*Carduelis chloris*) | count |
| Certhia.brachydactyla | Abundance of short-toed treecreeper (*Certhia brachydactyla*) | count |
| Columba.palumbus | Abundance of common wood pigeon (*Columba palumbus*) | count |
| Corvus.corone | Abundance of carrion crow (*Corvus corone*) | count |
| Cuculus.canorus | Abundance of common cuckoo (*Cuculus canorus*) | count |
| Cyanistes.caeruleus | Abundance of Eurasian blue tit (*Cyanistes caeruleus*) | count |
| Dendrocopos.majo | Abundance of great spotted woodpecker (*Dendrocopos major*) | count |
| Emberiza.citrinella | Abundance of yellowhammer (*Emberiza citrinella*) | count |
| Erithacus.rubecula | Abundance of European robin (*Erithacus rubecula*) | count |
| Fringilla.coelebs | Abundance of common chaffinch (*Fringilla coelebs*) | count |
| Garrulus.glandarius | Abundance of Eurasian jay (*Garrulus glandarius*) | count |
| Parus.major | Abundance of great tit (*Parus major*) | count |
| Passer.domesticus | Abundance of house sparrow (*Passer domesticus*) | count |
| Phoenicurus.ochruros | Abundance of black redstart (*Phoenicurus ochruros*) | count |
| Phylloscopus.collybita | Abundance of common chiffchaff (*Phylloscopus collybita*) | count |
| Pica.pica | Abundance of Eurasian magpie (*Pica pica*) | count |
| Picus.viridis | Abundance of European green woodpecker (*Picus viridis*) | count |
| Prunella.modularis | Abundance of dunnock (*Prunella modularis*) | count |
| Sitta.europaea | Abundance of Eurasian nuthatch (*Sitta europaea*) | count |
| Streptopelia.decaocto | Abundance of Eurasian collared dove (*Streptopelia decaocto*) | count |
| Streptopelia.turtur | Abundance of European turtle dove (*Streptopelia turtur*) | count |
| *Sturnus.vulgaris* | Abundance of common starling (*Sturnus vulgaris*) | count |
| *Sylvia.atricapilla* | Abundance of Eurasian blackcap (*Sylvia atricapilla*) | count |
| *Sylvia.communis* | Abundance of common whitethroat (*Sylvia communis*) | count |
| *Troglodytes.troglodytes* | Abundance of Eurasian wren (*Troglodytes troglodytes*) | count |
| *Turdus.merul* | Abundance of common blackbird (*Turdus merula*) | count |
| *Turdus.philomelos* | Abundance of song thrush (*Turdus philomelos*) | count |

##### LUCAS

All variables of this dataset, their definitions and sources are listed in Table 6. Climate variables were obtained from CHELSA V2.1 at 1 km resolution for 1981–2010 [48, 15]. Topsoil physical and chemical properties were obtained from the in-situ measurements of the LUCAS Soil surveys of 2009, 2015 and 2018 [93, 57, 45, 28], except available water capacity, which was taken from the LUCAS-based topsoil physical properties for Europe [10]. Heavy metal concentrations were obtained from the LUCAS topsoil heavy metals dataset [92], and soil loss by water erosion from [60]. Geological variables were extracted from the European Soil Database v2 1 km raster library [25]. All soil datasets are distributed by the European Soil Data Centre [62, 61] at https://esdac.jrc.ec.europa.eu/, subject to its data licence agreement. Topographic variables were derived from the Copernicus DEM [27] in ArcGIS Pro [24], with the terrain ruggedness index following [73]. Hydrographic distances were computed in ArcGIS Pro from the EU-Hydro river network and coastal polygons [26].

**Table 6:**
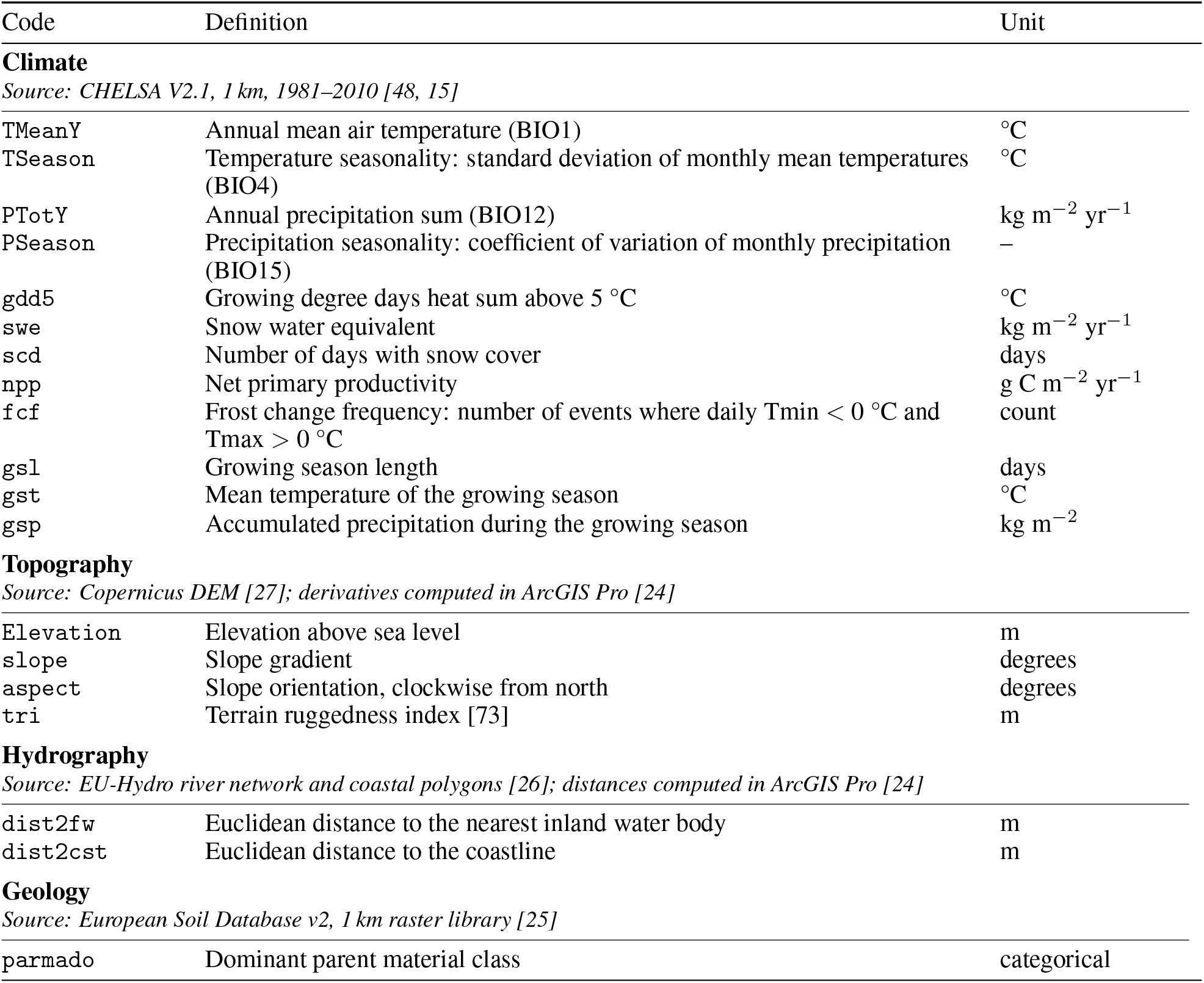
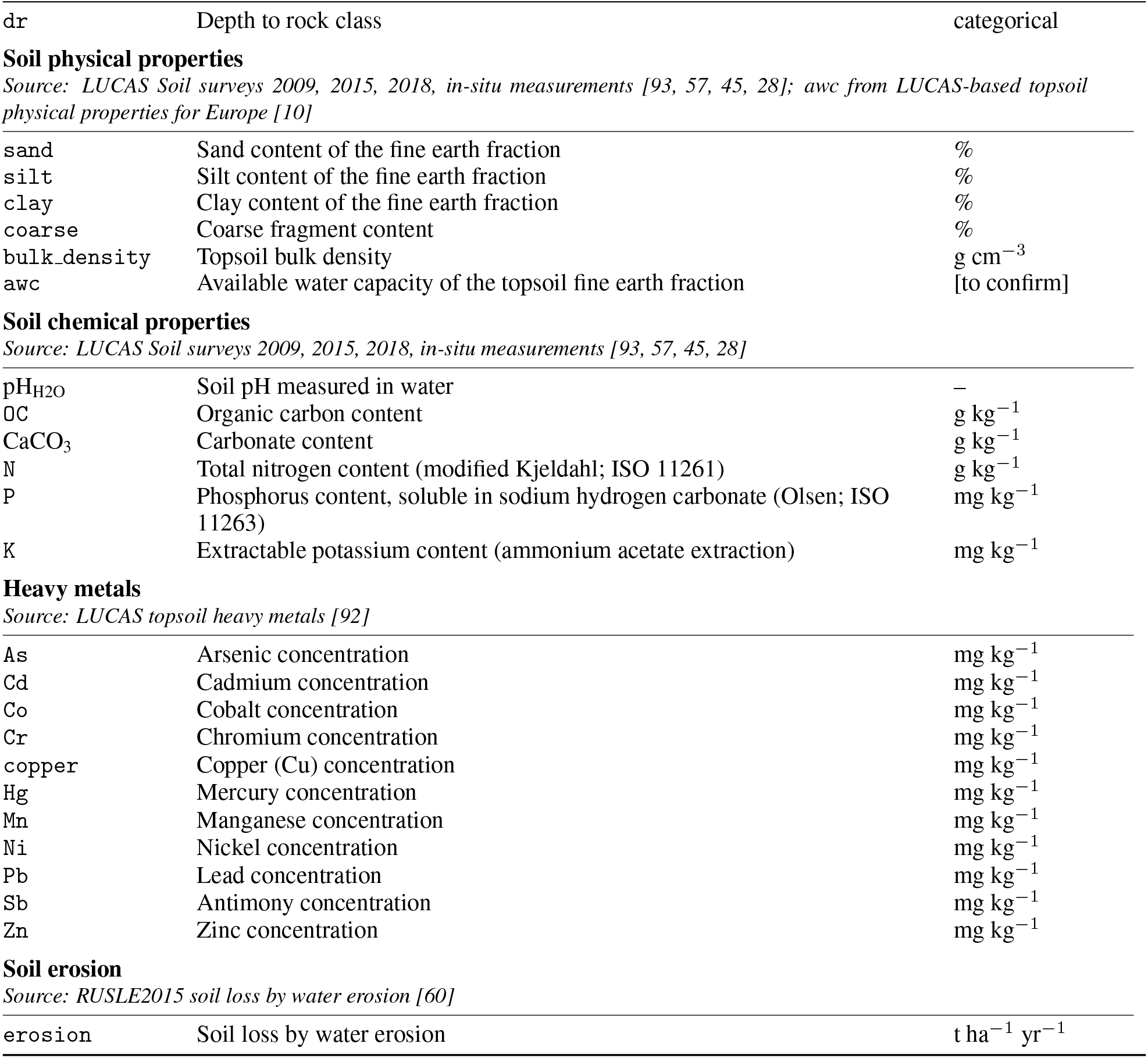
Variable definitions and data sources, grouped by feature type for the LUCAS dataset.

| Code | Definition | Unit |
| --- | --- | --- |
| **Climate** |  |  |
| *Source: CHELSA V2.1, 1 km, 1981–2010 [48, 15]* |  |  |
| TMeanY | Annual mean air temperature (BIO1) | °C |
| TSeason | Temperature seasonality: standard deviation of monthly mean temperatures (BIO4) | °C |
| PTotY | Annual precipitation sum (BIO12) | kg m<sup>-2</sup> yr<sup>-1</sup> |
| PSeason | Precipitation seasonality: coefficient of variation of monthly precipitation (BIO15) | – |
| gdd5 | Growing degree days heat sum above 5 °C | °C |
| swe | Snow water equivalent | kg m<sup>-2</sup> yr<sup>-1</sup> |
| scd | Number of days with snow cover | days |
| npp | Net primary productivity | g C m<sup>-2</sup> yr<sup>-1</sup> |
| fcf | Frost change frequency: number of events where daily Tmin < 0 °C and Tmax > 0 °C | count |
| gsl | Growing season length | days |
| gst | Mean temperature of the growing season | °C |
| gsp | Accumulated precipitation during the growing season | kg m<sup>-2</sup> |
| **Topography** |  |  |
| *Source: Copernicus DEM [27]; derivatives computed in ArcGIS Pro [24]* |  |  |
| Elevation | Elevation above sea level | m |
| slope | Slope gradient | degrees |
| aspect | Slope orientation, clockwise from north | degrees |
| tri | Terrain ruggedness index [73] | m |
| **Hydrography** |  |  |
| *Source: EU-Hydro river network and coastal polygons [26]; distances computed in ArcGIS Pro [24]* |  |  |
| dist2fw | Euclidean distance to the nearest inland water body | m |
| dist2cst | Euclidean distance to the coastline | m |
| **Geology** |  |  |
| *Source: European Soil Database v2, 1 km raster library [25]* |  |  |
| parmado | Dominant parent material class | categorical |
| dr | Depth to rock class | categorical |
| **Soil physical properties** |  |  |
| *Source: LUCAS Soil surveys 2009, 2015, 2018, in-situ measurements [93, 57, 45, 28]; awc from LUCAS-based topsoil physical properties for Europe [10]* |  |  |
| sand | Sand content of the fine earth fraction | % |
| silt | Silt content of the fine earth fraction | % |
| clay | Clay content of the fine earth fraction | % |
| coarse | Coarse fragment content | % |
| bulk_density | Topsoil bulk density | g cm<sup>-3</sup> |
| awc | Available water capacity of the topsoil fine earth fraction | [to confirm] |
| **Soil chemical properties** |  |  |
| *Source: LUCAS Soil surveys 2009, 2015, 2018, in-situ measurements [93, 57, 45, 28]* |  |  |
| pH<sub>H2O</sub> | Soil pH measured in water | – |
| OC | Organic carbon content | g kg<sup>-1</sup> |
| CaCO<sub>3</sub> | Carbonate content | g kg<sup>-1</sup> |
| N | Total nitrogen content (modified Kjeldahl; ISO 11261) | g kg<sup>-1</sup> |
| P | Phosphorus content, soluble in sodium hydrogen carbonate (Olsen; ISO 11263) | mg kg<sup>-1</sup> |
| K | Extractable potassium content (ammonium acetate extraction) | mg kg<sup>-1</sup> |
| **Heavy metals** |  |  |
| *Source: LUCAS topsoil heavy metals [92]* |  |  |
| As | Arsenic concentration | mg kg<sup>-1</sup> |
| Cd | Cadmium concentration | mg kg<sup>-1</sup> |
| Co | Cobalt concentration | mg kg<sup>-1</sup> |
| Cr | Chromium concentration | mg kg<sup>-1</sup> |
| copper | Copper (Cu) concentration | mg kg<sup>-1</sup> |
| Hg | Mercury concentration | mg kg<sup>-1</sup> |
| Mn | Manganese concentration | mg kg<sup>-1</sup> |
| Ni | Nickel concentration | mg kg<sup>-1</sup> |
| Pb | Lead concentration | mg kg<sup>-1</sup> |
| Sb | Antimony concentration | mg kg<sup>-1</sup> |
| Zn | Zinc concentration | mg kg<sup>-1</sup> |
| **Soil erosion** |  |  |
| *Source: RUSLE2015 soil loss by water erosion [60]* |  |  |
| erosion | Soil loss by water erosion | t ha<sup>-1</sup> yr<sup>-1</sup> |

#### G.2 Main results

##### STOC

In the following, we give the partial causal orders given by PC and by RestPC. Results for the partial causal order obtained by PC:

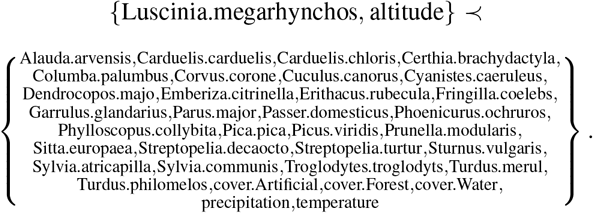

Results for the partial causal order obtained by RestPC:

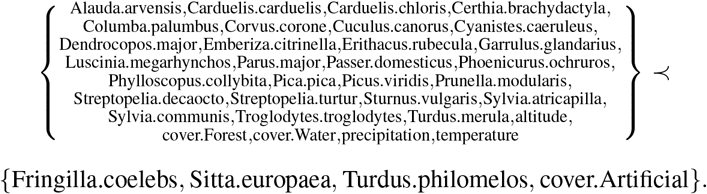

##### LUCAS

In the following, we give the partial causal orders of PC and RestPC. Partial causal order obtained using the PC algorithm:

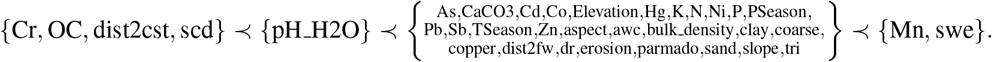

Partial causal order obtained using the RestPC algorithm:

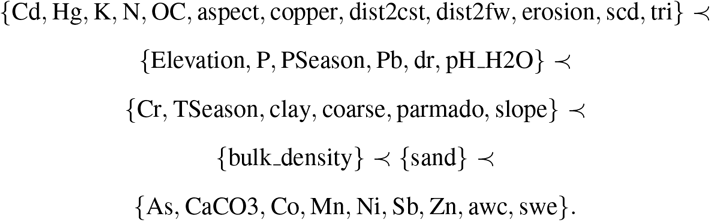

#### G.3 Sensitivity analysis

##### STOC

We conducted a sensitivity analysis using different values of *β* = [0.4, 0.5, 0.6], where *β* = 0.5 was used in the main analysis. Larger values of *β* lead to a sparser graph. A summary of the results is presented in Table 7 for PC and in Table 8 for RestPC. In each table, the columns correspond to the different values of *β*, while *L*_*i*_ denotes assignments of the variables associated with species abundance or environmental characteristics to partial causal order level *i*. Across both methods, the maximum number of levels remained two. For PC, the results were more sensitive to the value of *β*. At *β* = 0.4, PC identified a single partial causal order level, whereas for larger values of *β* the variables were separated into two partial causal order levels. Only `Luscinia.megarhynchos` and altitude have the same partial causal order across all values of *β*. Interestingly, as in RestPC, `temperature` was assigned to *L*_2_ when *β* = 0.6. For RestPC, the overall two-level structure was preserved across the different values of *β*, although some variables changed partial order level. In particular, compared with the main analysis at *β* = 0.5, the assignments of `Sitta.europaea`, `Luscinia.megarhynchos`, and temperature changed when *β* = 0.6. Nevertheless, most variables remained in the same partial causal order level, suggesting that the partial causal ordering obtained by RestPC is relatively stable with respect to the choice of *β*. One possible interpretation is that the species assigned to *L*_1_ are not directly associated with temperature, whereas variables or species assigned to *L*_2_ may be more closely related to temperature-dependent processes. Since altitude can influence temperature, part of the apparent relationship between temperature and species abundance may be explained by altitude.

**Table 7:** Level membership in the partial causal order obtained by PC for different values of *β*.

| Variable | $\beta = 0.4$ | $\beta = 0.5$ | $\beta = 0.6$ |
| --- | --- | --- | --- |
| `Luscinia.megarhynchos` | $L_1$ | $L_1$ | $L_1$ |
| `altitude` | $L_1$ | $L_1$ | $L_1$ |
| `temperature` | $L_1$ | $L_2$ | $L_2$ |
| All remaining variables | $L_1$ | $L_2$ | $L_1$ |

For each value of $\beta$ , the partial causal ordering is $L_1 \prec L_2$ .

**Table 8:** Level membership in the partial causal order obtained by RestPC for different values of *β*.

| Variable | $\beta = 0.4$ | $\beta = 0.5$ | $\beta = 0.6$ |
| --- | --- | --- | --- |
| Fringilla.coelebs | $L_2$ | $L_2$ | $L_2$ |
| Sitta.europaea | $L_1$ | $L_2$ | $L_1$ |
| Turdus.philomelos | $L_1$ | $L_2$ | $L_2$ |
| cover.Artificial | $L_1$ | $L_2$ | $L_2$ |
| Luscinia.megarhynchos | $L_1$ | $L_1$ | $L_2$ |
| temperature | $L_1$ | $L_1$ | $L_2$ |
| All remaining variables | $L_1$ | $L_1$ | $L_1$ |

For each value of $\beta$ , the partial causal ordering is $L_1 \prec L_2$ .

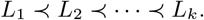

##### LUCAS

Similarly to the STOC dataset, we conducted a sensitivity analysis for different values of *β*. A summary of the results is presented in Table 9 for PC and in Table 10 for RestPC. For RestPC, the number of clusters ranged from four to six across the different values of *β*. Two sets of variables consistently occupied the first and last positions in the partial causal ordering, respectively: `Cd, Hg, K, N, OC,aspect, copper, dist2cst, dist2fw, scd`, and `tri` in the first level, and `As, CaCO3, Co, Mn, Ni,Sb, Zn, awc`, and `swe` in the last level. Overall, the RestPC results were relatively stable across values of *β*, with most changes occurring in the composition of the intermediate levels. For PC, the number of partial causal order levels ranged from three to eight, indicating greater sensitivity to the choice of *β*. Only a few variables were consistently assigned to the first or last level: `Cr` and `dist2cst` remained in *L*_1_, while Mn consistently appeared in the last level. The remaining variables showed more variation in their relative positions across values of *β*, suggesting that the PC-based partial causal ordering was less stable than the RestPC ordering.

**Table 9:**
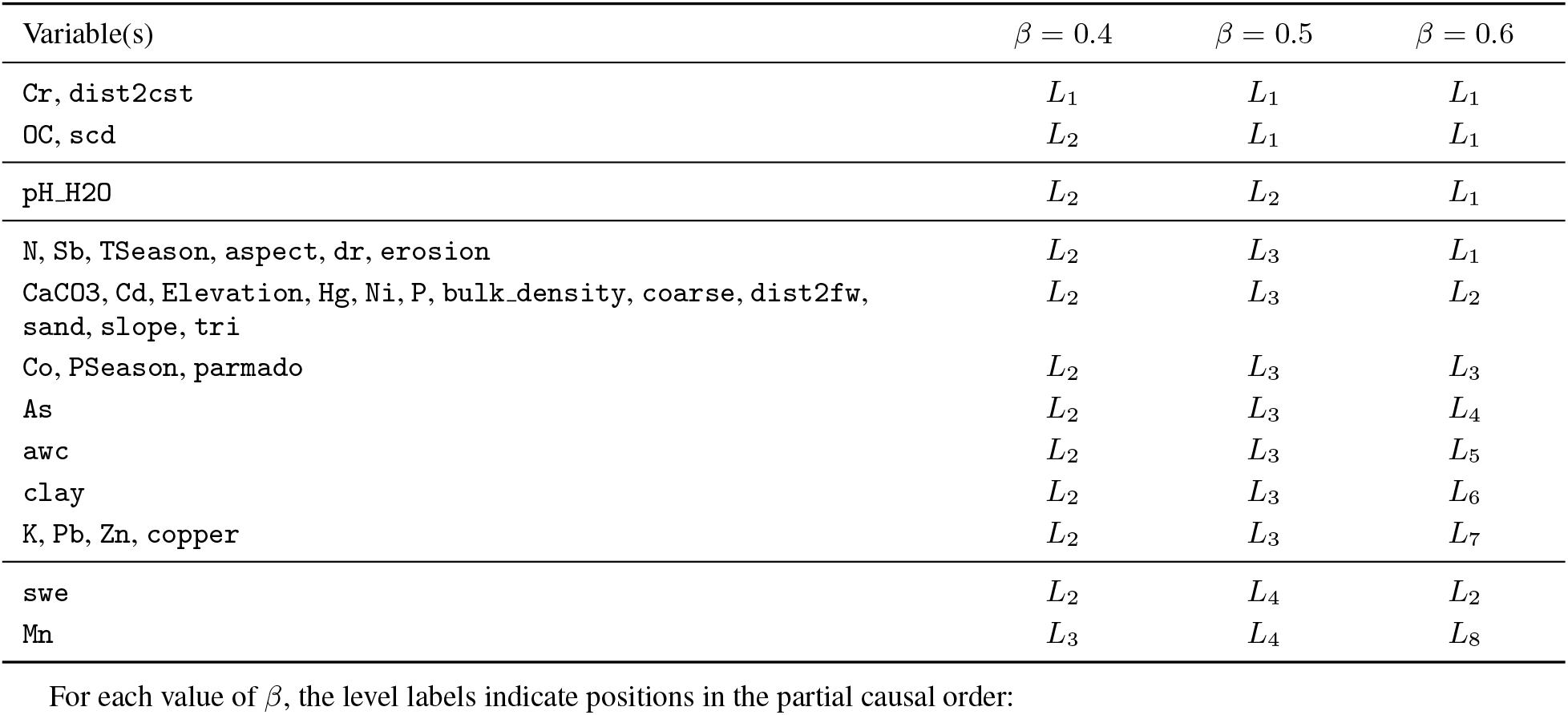
Level membership in the partial causal order obtained by PC for different values of *β*.

**Table 10:** Level membership in the partial causal order obtained by RestPC for different values of *β*.

| Variable(s) | $\beta = 0.4$ | $\beta = 0.5$ | $\beta = 0.6$ |
| --- | --- | --- | --- |
| Cd, Hg, K, N, OC, aspect, copper, dist2cst, dist2fw, scd, tri | $L_1$ | $L_1$ | $L_1$ |
| erosion | $L_1$ | $L_1$ | $L_2$ |
| Elevation, P, PSeason | $L_1$ | $L_2$ | $L_2$ |
| dr | $L_2$ | $L_2$ | $L_2$ |
| Pb | $L_2$ | $L_2$ | $L_4$ |
| pH_H2O | $L_2$ | $L_2$ | $L_3$ |
| Cr, coarse | $L_2$ | $L_3$ | $L_5$ |
| TSeason, clay | $L_2$ | $L_3$ | $L_3$ |
| parmado | $L_2$ | $L_3$ | $L_6$ |
| bulk_density | $L_2$ | $L_4$ | $L_3$ |
| slope | $L_3$ | $L_3$ | $L_5$ |
| sand | $L_3$ | $L_5$ | $L_6$ |
| As, CaCO3, Co, Mn, Ni, Sb, Zn, awc, swe | $L_4$ | $L_6$ | $L_6$ |

For each value of $\beta$ , the level labels indicate positions in the partial causal order:

#### G.4 Negative controls

In addition to sensitivity analysis, we compare the partial causal orders produced by PC and RestPC with the negative controls adapting the approach proposed by [67]. For the metrics that can be expressed as a function of adjacency confusion matrices, the authors provide exact null distributions, using the maximum number of edges in the graph, the number of edges in the ground truth graph, and the number of edges in the estimated graphs. These distributions are used to compute the expected value under the null hypothesis for these metrics and introduce a statistical test for skeleton fit. For other metrics, the authors propose to use simulation-based negative controls, i.e to generate random

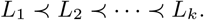

negative control graphs that match relevant characteristics of the algorithmic estimate, and evaluate the algorithmic performance with the corresponding empirical negative control distribution. In this case, for a statistical comparison, the authors suggest using the empirical p-value as:

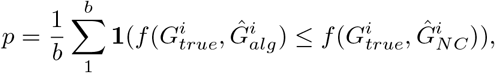

where *b* is the number of generated random control graphs, *f* denotes a performance measure for which smaller values indicate better performance (otherwise, the inequality is reversed). In our study, we do not aim to evaluate the graph obtained by the algorithm, but we evaluate the produced partial causal orders. Because neither a ground-truth graph nor a complete causal order is available, we evaluate the results against a limited set of pre-specified causal-order relationships. We therefore adapt the negative-control approach of [67] to this setting.

For both datasets, we applied PC and RestPC and compared the estimated partial causal orders with the pre-specified causal-order relationships. Both algorithms produced partial orders in which many of these relationships remained unresolved. To construct comparable negative controls, we aimed to preserve the structural resolution of each algorithmic estimate. For each dataset and algorithm, we generated *b* = 10000 negative-control orders by randomly permuting the variable labels while preserving the number, sizes, and ordering of the estimated clusters. This conditional permutation preserves the overall resolution implied by the cluster structure, that is, the total number of all possible variable pairs that the structure can resolve, while the number of pre-specified relationships classified as unresolved may vary across permutations. We then tested whether the observed alignment with the pre-specified relationships was stronger than expected under random label assignment. We evaluate the alignment using the numbers of correctly ordered, reversed and unresolved relationships, with empirical p-values calculated for each measure [68, 67].

For STOC dataset, the results of the negative control analysis are presented in Table 11. On this dataset, the PC algorithm has more correct and fewer reversed relationships than the corresponding expected mean under the empirical null distribution. RestPC showed the opposite pattern, with fewer correct and more reversed relationships than expected under the negative control, although RestPC has slightly fewer unresolved relationships than expected on average. None of these comparisons reached the 5% significance threshold, with the smallest p-value p =0.07 for the number of correct and unresolved relationships produced by PC.

**Table 11:** Comparison of the observed causal-order performance of PC and RestPC with the permutation-based negative control on STOC.

| Method | Metric | Observed | Null mean | 95% null interval | Permutation $p\text{-value}$ |
| --- | --- | --- | --- | --- | --- |
| **PC** | Correct | 31 | 9.6 | [0, 31] | 0.07 |
|  | Reversed | 5 | 9.9 | [5, 12] | 0.21 |
|  | Unresolved | 146 | 162.5 | [146, 170] | 0.07 |
| **RestPC** | Correct | 15 | 18.5 | [8, 24] | 0.88 |
|  | Reversed | 27 | 18.5 | [0, 58] | 0.81 |
|  | Unresolved | 140 | 145.1 | [116, 158] | 0.54 |

The results for the LUCAS dataset are presented in Table 12. On this dataset, the PC algorithm has fewer correct and fewer reversed relationships than the corresponding null means, which reflects a higher number of unresolved relationships than the corresponding null mean. RestPC produced more correct and fewer reversed relationships, while at the same time having a lower number of unresolved relationships in comparison to the expected mean under the negative control. RestPC yielded the smallest p-values for the numbers of correct and reversed relationships ((p=0.07) and (p=0.15), respectively), although neither reached the conventional 5% significance threshold. However, RestPC has a significantly lower number of unresolved relationships (p-value =0.04)

**Table 12:** Comparison of the observed causal-order performance of PC and RestPC with the permutation-based negative control on LUCAS.

| Method | Metric | Observed | Null mean | 95% null interval | Permutation $p\text{-value}$ |
| --- | --- | --- | --- | --- | --- |
| **PC** | Correct | 38 | 54.3 | [14, 97] | 0.78 |
|  | Reversed | 49 | 54.2 | [18, 92] | 0.42 |
|  | Unresolved | 220 | 198.6 | [177, 221] | 0.97 |
| **RestPC** | Correct | 159 | 119.2 | [71, 174] | 0.07 |
|  | Reversed | 91 | 119.7 | [69, 174] | 0.15 |
|  | Unresolved | 57 | 68.1 | [56, 77] | 0.04 |

Overall, neither PC nor RestPC produces a significantly larger number of correctly ordered relationships than its permutation-based negative control at the 5% level. Nevertheless, PC on STOC and RestPC on LUCAS both yield more correctly ordered relationships than expected on average under the null, with a p-value= 0.07 in both cases. Moreover, on LUCAS, RestPC produces significantly fewer unresolved relationships than expected under the negative control with a p-value= 0.04. Our procedure is a heuristic adaptation of [67] to partial causal orders. The results should therefore be interpreted as exploratory evidence rather than as formal validation.

### H Micro causal relations and single-time-point data

In this section, we use the same examples as in the main manuscript (in particular, Figure 1 of the main manuscript) to illustrate why micro-level causal relations cannot be recovered from single-time-point data.

#### Micro-level relations

When only variables at time *t* are observed, the PC algorithm cannot in general recover the correct skeleton among the variables inV_*t*_. Nevertheless, it can still reveal the absence of some relations. For instance, if two variables *X*_*t*_ and *Y*_*t*_ belong to different components of the full-time DAG^2^, then they are necessarily marginally independent, since no path connects them. More generally, when two variables belong to the same component, the PC algorithm can detect the absence of a relation between them only if they are already marginally independent. This is illustrated in Figure 1(c,g,k,o) in the main text, which shows the outputs of the PC algorithm obtained from data generated respectively from the full-time DAGs in Figure 1(a,e,i,m) in the main text. In panels (c) and (g), the PC algorithm is unable to discover that there is no relation between *X*_*t*_ and *Y*_*t*_, because establishing their independence would require conditioning on past variables, which are unobserved. By contrast, in panels (k) and (o), the algorithm correctly detects the absence of a relation between *X*_*t*_ and *Y*_*t*_, since these variables are already marginally independent.

Orientation is even more problematic. From observations restricted to time *t*, one cannot distinguish between two fundamentally different situations. In the first, *X*_*t*_ directly causes *Y*_*t*_. In the second, the dependence between *X*_*t*_ and *Y*_*t*_ is induced by an unobserved past variable. For example, suppose that *X*_*t*−1_ causes *Y*_*t*_. Under Assumption 2.4, *X*_*t*−1_ also causes *X*_*t*_, so that *X*_*t*−1_ becomes an unobserved common cause of *X*_*t*_ and *Y*_*t*_. As a result, both scenarios produce the same qualitative pattern in the observed data: *X*_*t*_ and *Y*_*t*_ remain dependent given any subset of the observed variables inV_*t*_, and single-time-point data alone do not allow one to determine whether the dependence between *X*_*t*_ and *Y*_*t*_ reflects a genuine micro-level causal relation at time *t*, or is instead induced by the unobserved past. We therefore conclude that micro-level causal relations between variables inV_*t*_ are, in general, not identifiable from such data. Consistent with this, in Figure 1(c,g,k,o) in the main text, all edge orientations returned by the PC algorithm among variables inV_*t*_ do not correspond to the true micro-level causal relations.

One may argue that, in this setting, the FCI algorithm is theoretically more appropriate than PC, since it explicitly allows for unmeasured confounding. However, because hidden temporal connections are pervasive in the present context, FCI is unlikely to provide substantially more information than PC beyond the overall dependence structure. For example, the outputs of the FCI algorithm shown in Figure 1(d,h,l,p) are no more informative than those of the PC algorithm in these examples.

Overall, under our assumptions, micro-level causal relations cannot in general be recovered from data observed at a single time point. Such data can at best be used to infer the absence of some simple relations. This is already well established in the literature [36, 50, 79, 77].

### I Illustration of single-time point data

In Figure 7 we provide an illustration of the generation process underlying the single-time-point data. In case of complete observability, we assume that the data is generated from a dynamical system that evolves over time and can therefore be represented as a time series. Furthermore, we observe this system in multiple locations. However, we consider that complete observability is not satisfied, we only have access to a single time point in each location. Specifically, the same system is observed across multiple locations at a common sampling time *t*, which is assumed to occur long after the onset of the dynamics. The underlying time series *X, Y*, and *Z* in Figure 7 may represent species, functions, services, or environmental factors. On the right of Figure 7, each window labelled Sample 1, …, Sample *n*, corresponds to a different location, while the vertical line indicates the time point at which that location is actually observed. As shown in the right panel of Figure 7, these single-time-point observations collectively form a cross-sectional dataset, in which each sample corresponds to a distinct location and no explicit temporal dimension is retained.

**Figure 7.**
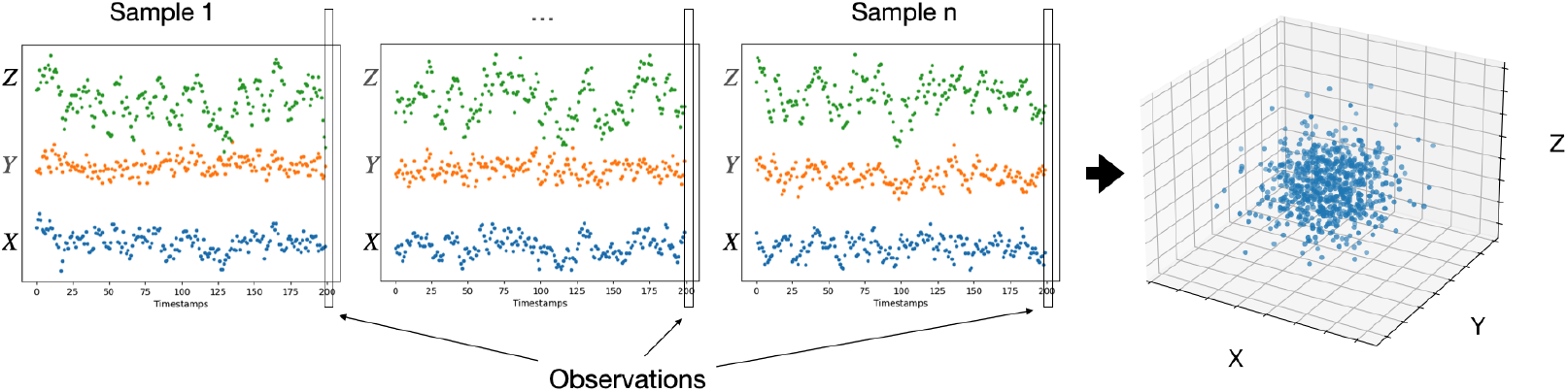
Illustration of ecological data consisting of *n* observations and 200 time-points of three processes *X, Y*, and *Z*. Left: time-series observations across multiple sampling locations. Right: single-time-point observations extracted from the last time-point. For simplicity, time is assumed to be aligned across locations but this is not a requirement for the validity of our results.

### J Data accessibility statement

- The processed STOC dataset used in this study were obtained from [31]. Minor modifications to the original preprocessing procedure are described in Section G.1. In accordance with the data availability statement of that publication, the data and associated preprocessing code are publicly available in the Zenodo repository at https://doi.org/10.5281/zenodo.10229534.
- The processed LUCAS dataset is publicly available in the Zenodo repository at https://doi.org/10.5281/zenodo.22845926.

### Footnotes

1 The acyclicity assumption is common but may be violated in ecology, for instance through instantaneous feedback in food webs [95]. More generally, acyclicity remains difficult to assess empirically, even though a common view is that cyclicity arises mainly at the process level rather than in the underlying full-time graph [74].

2 A component of a graph is a connected subgraph that is not contained in any larger connected subgraph.

